# The neural circuit implementation of distributional reinforcement learning

**DOI:** 10.64898/2026.11.07.765111

**Authors:** Sara Matias, Camille Rullán Buxó, Shudi Xu, Malcolm G. Campbell, Adam S. Lowet, Xiaotang Lu, Yuelong Wu, Richard Schalek, Jeff W. Lichtman, Mitsuko Watabe-Uchida, Jan Drugowitsch, Naoshige Uchida

**Affiliations:** Center for Brain Science, Harvard University, Cambridge, Massachusetts, USA; Department of Molecular and Cellular Biology, Harvard University, Cambridge, Massachusetts, USA; Department of Neurobiology, Harvard Medical School, Boston, Massachusetts, USA; Interdepartmental Neuroscience, Northwestern University, Chicago, Illinois, USA; Li Ka Shing Center, University of California Berkeley, Berkeley, California, USA; Department of Chemistry, University of Illinois Urbana-Champaign, Urbana, Illinois, USA; Kempner Institute for the Study of Natural & Artificial Intelligence, Harvard University, Allston, Massachusetts, USA

**Keywords:** Distributional reinforcement learning, hierarchical dopamine projections, striatum, distributional actor-critic, optimism spectrum

## Abstract

Distributional reinforcement learning (RL) algorithms outperform traditional RL by representing the full probability distribution of future outcomes rather than its mean^1^. Neural signatures consistent with distributional coding have been observed in dopamine neurons^2^ and in striatal value representations^3^. However, how distributional RL is implemented within the anatomical circuit organization of the basal ganglia remains unknown. Uncovering how representations of outcome distributions are organized across brain circuits has broad implications for both fundamental and clinical neuroscience, providing new insights into how the brain represents uncertainty and how these representations shape risk-sensitive learning and decision-making^4–6^. Here, we investigated how representations of outcome distributions are organized across dopamine projections to the striatum by combining striatum-wide fiber photometry of dopamine axonal activity with Neuropixels recordings from projection-defined dopamine neurons. We found that distributional reward prediction error signals are preferentially encoded in lateral ventral striatum-projecting dopamine neurons, but that reward distribution information is shared between the ventral and the dorsal striatum. Additionally, an anatomical gradient of optimism in the midbrain supports the diversity of value predictions required for distributional RL. Our findings extend the actor–critic framework proposed for the basal ganglia^7–9^ by revealing distributional prediction errors in a subpopulation of mesolimbic dopamine neurons capable of training parallel striatal regions thought to implement different types of behavioral control^10–14^.

## 1 Introduction

The long-standing dialogue between research on natural and artificial intelligence has inspired the development of reinforcement learning (RL), now one of the most influential frameworks for studying learning and decision-making, in which agents learn to maximize the rewards they obtain in an environment. Many RL algorithms learn by minimizing prediction errors, which are defined as the difference between actual and expected rewards. Minimizing this simple cost function yields remarkably useful value representations that support adaptive behaviors^15–20^. Nevertheless, reducing the prediction of future outcomes to a single scalar value inevitably discards information about their uncertainty and structure. *Distributional* reinforcement learning addresses this limitation by representing the probability distribution of future outcomes^1,21^.

Distributional agents outperform traditional ones across multiple benchmarks, including complex video games and continuous control problems^1,2,22–25^. Rather than simply retaining more detail, distributional RL preserves information that is directly relevant for decision-making by representing the range and likelihood of possible future outcomes. This richer representation allows agents to flexibly adjust behavioral policies, for example, by emphasizing the lower or upper portions of the return distribution to generate risk-averse or risk-seeking decisions^1,26–29^.

In the brain, RL frameworks have provided an influential account of learning, with phasic activity of dopamine neurons encoding reward prediction error (RPE) signals of temporal difference learning algorithms^18,30–34^. Using dopamine teaching signals, the striatum, in turn, learns value representations and behavioral policies^3,34,35^. Remarkably, recent studies have shown that the activity of dopamine neuron populations^2^, as well as their striatal targets^3^, exhibit signatures of distributional RL.

Altogether, distributional RL offers a simple and biologically plausible framework for representing entire outcome distributions^2,3,23,36^, yet how the brain implements it remains poorly understood. Uncovering the underlying circuitry could advance our understanding of risk-sensitive decision-making and potentially enable its manipulation^37^. Indeed, choice behavior depends not only on expected outcomes but also on their probability distributions and the decision-maker’s risk preferences^38–40^. Dysregulation of these computations could contribute to maladaptive decision-making^41^. Several mechanisms have been proposed for how the basal ganglia might represent outcome uncertainty, including in the sustained activity of dopamine neurons^42,43^ (but see^30^), or in the sum of synaptic weights of striatal neurons expressing the D1 and D2 dopamine receptors^44^. However, these proposals are not derived from a normative theory of learning nor do they provide a conceptual framework to investigate how the dopamine circuitry is anatomically organized to implement them. Distributional RL is different: it represents more properties of the distribution, beyond its average and variance, and it makes specific predictions about the anatomical organization of dopamine neurons and their projections to the striatum to support its computational principle. Here, we sought to uncover the neural circuit organization supporting distributional RL in the basal ganglia using multi-site fiber photometry and projection-specific single unit recordings of dopamine neurons. By spatially mapping dopamine signals across the striatum and the midbrain, we found that the reward distribution is represented in dopamine populations projecting to the lateral ventral striatum and that these information are shared with the dorsal striatum. Although these striatal regions receive dopamine signals spanning the entire reward distribution, we found that neurons encoding its lower and upper tails are organized along an anatomical gradient in the midbrain, consistent with a circuit architecture that preserves distinct distributional value predictions^2^.

## 2 Results

### 2.1 Anatomical predictions from the distributional RL framework

Distributional RL offers several methods for representing reward distributions^1,21,45^, but experimental tests of distributional coding in the brain have focused on expectiles, since they naturally extend the learning rule used in traditional RL^23,46^. Just as quantiles generalize the median, expectiles generalize the mean^45^. Together, a family of expectiles characterizes the shape of a distribution, distinguishing distributions with identical means (Fig. 1a,b). Each expectile, 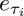, is obtained by minimizing an asymmetrically weighted squared-error loss function^23^ characterized by its asymmetry, *τ_i_* (Fig. 1b). This minimization corresponds, in the brain, to the learning process of minimizing the prediction error term from an asymmetrically weighted value update. Thus, distributional RL assumes a diverse population of value predictors, *V_i_*, each updated with its own pair of learning rates, 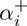 and 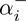, for positive and negative prediction errors, respectively^2^. When the two learning rates are equal, *α*^+^ = *α^−^*, as in traditional RL, every predictor converges to the mean of the reward distribution (Fig. 1c). When they differ, learning is biased: predictors with *α*^+^ *< α^−^* converge below the mean and are pessimistic, whereas predictors with *α*^+^ *> α^−^* converge above it and are optimistic (Fig. 1d). We quantify this asymmetry with an optimism index, 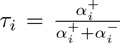, which ranges from 0 (fully pessimistic) to 1 (fully optimistic).

**Fig. 1.**
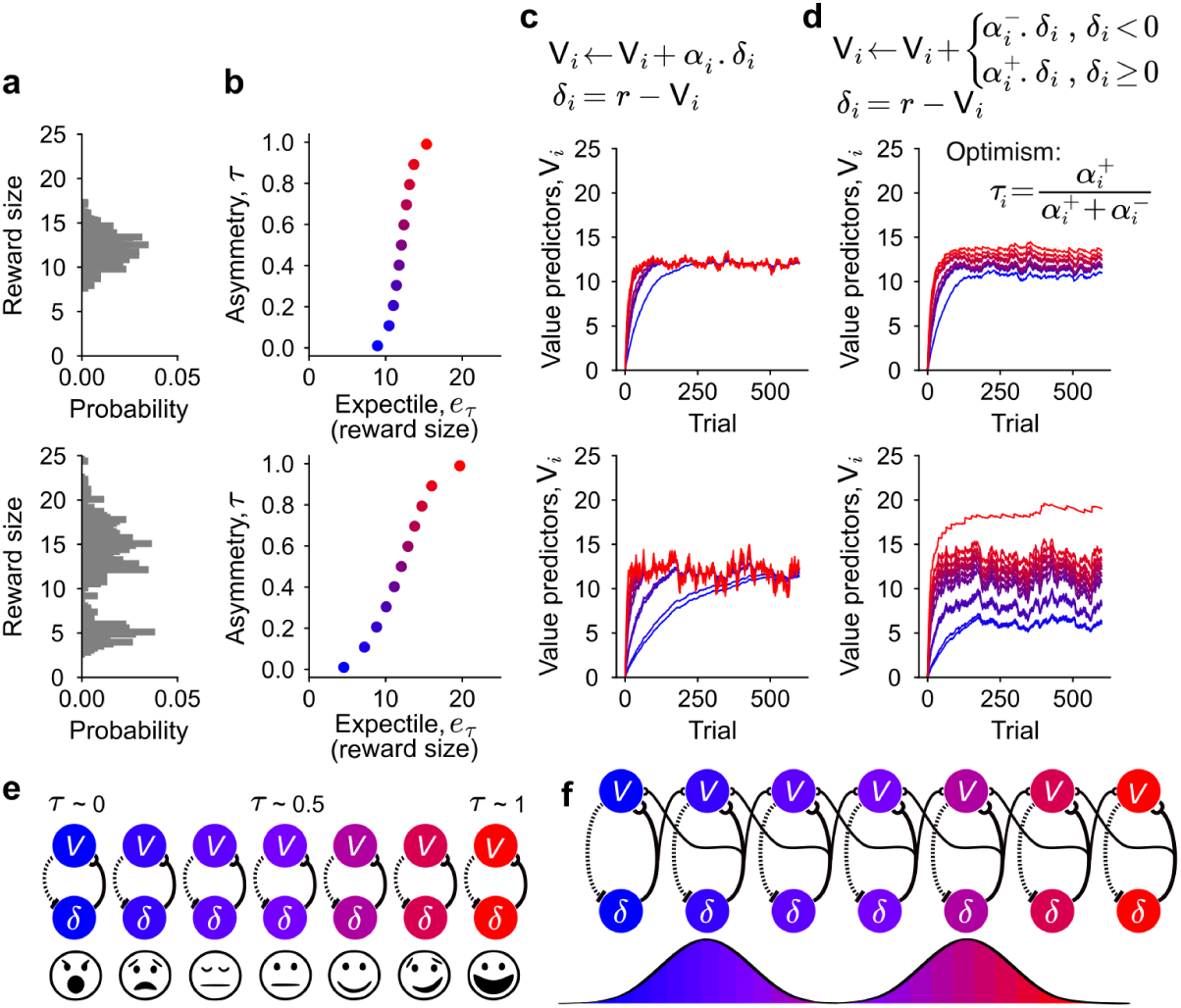
Connectivity required by distributional RL. **a,** Two distributions with the same mean but different shape (top and bottom) can be distinguished using their expectiles. **b,** Inverse expectile function for each one of the distributions in **a**, which is unique to each distribution. **c,** Diversity in learning rates in the absence of asymmetric scaling cause value predictors (*V_i_* in colors) to converge asymptotically to a single scalar value: the expected value of the distribution. **d,** Diversity in positive and negative learning rates (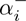 and 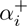) leads to convergence at different reward sizes for different value predictors. The optimism level, *τ*, is derived from the relationship between the learning rates to positive and negative prediction errors: pessimism corresponds to an optimism level close to zero, and high optimism close to one. **e,** The simplest implementation of a loop structure between RPEs, *δ_i_*, and value predictors, *V_i_*, would be a one-to-one mapping between them, but such architecture is biologically implausible. **f**, A Gaussian-weighted topographic connectivity between RPEs and value predictors has been proposed as a solution that enables distributional RL^2^, such that cross-talk between loops decrease with increasing optimism distance.

Because prediction errors are calculated using value predictions, diversity in the population of value predictors causes diversity in the population of prediction errors. In fact, the reward magnitude at which the prediction error is zero (*δ_i_* = 0), termed the zero-crossing point, is the reward for which *V_i_* receives no update^2,23^. Each prediction error neuron is, therefore, characterized by two parameters: its optimism, an intrinsic property of the neuron, and its zero crossing point, which shifts with the distribution being learned. Because more optimistic predictors tend to converge to higher expectiles, the two parameters must always be positively correlated across the population (Fig. 1b); this is the key signature of distributional RL which we test below.

Mapped onto the brain, striatal neurons represent the value predictors^3^, *V_i_*, dopamine neurons compute RPEs, *δ_i_*, and dopamine release in turn updates striatal value representations *V_i_* through a recurrent connectivity loop^34^ (Fig. 1e). The learning rate parameters, *α*^+^ and *α^−^*, are correlated with the slopes of the piecewise linear response function of dopamine neurons to positive and to negative prediction errors, respectively^2^(Fig. 1d). For this learning loop to work, optimistic and pessimistic updates must be kept separate, so that each value predictor is updated by RPEs with matching asymmetry. The dense and widespread axonal arborizations of dopamine neurons in the nucleus accumbens (NAc)^47^ and in the dorsal striatum (DS)^48^, however, appears incompatible with a specific, one-to-one loop between midbrain dopamine neurons and striatal neurons. One proposed solution is that dopamine-striatal connectivity is organized along an anatomical gradient of optimism that preserves biased value updates^2,49^(Fig. 1f). Another is that separation arises from the distinct dopamine signaling properties of striatal cell types^3^. It also remains unclear how any such structured RPE heterogeneity coexists with the functional diversity of dopamine responses observed across the striatum, including signals related to environmental and internal variables beyond canonical RPEs^50–52^.

To address this question, we characterized axonal and cell body dopamine activity projecting to multiple striatal areas using projection-specific approaches.

### 2.2 Anatomical gradient of canonical RPEs across the striatum

RPE-like activity has been observed in both the ventral and the dorsal striatum in simple Pavlovian and operant tasks^34,49,53^. However, dopamine signals differ across striatal regions^50,51,54–56^ and very few studies have systematically mapped both positive and negative prediction error signals simultaneously across the entire striatum^49,57^. Because learning from both positive and negative RPEs is essential for convergence of value learning in both traditional and distributional RL, we first asked which striatal regions receive relatively intact, bi-directional (i.e. positive and negative) RPE signals at the population level, which we will refer to as “canonical RPEs” (irrespective of asymmetry).

We used multi-fiber photometry to record calcium activity from dopamine axons across multiple striatal regions using a custom implant containing fourteen optical fibers. Recordings were performed in triple-transgenic DAT-Cre::Ai148::Ai14 mice expressing the calcium indicator GCaMP6f^58^ together with tdTomato^59^, which served as an activity-independent control signal, in dopamine neurons (Fig. 2b). Optical fibers targeted the medial nucleus accumbens (mNAc), the ventral nucleus accumbens (vNAc, *×*2), the lateral nucleus accumbens (lNAc), the dorsomedial striatum (DMS, *×*4), the dorsolateral striatum (DLS, *×*4), and the tail of the striatum (TS, *×*2). We used two CMOS cameras to record isosbestic, green and red fluorescence from all optical fibers simultaneously (Extended Data Fig. 1) and post-mortem micro-computed tomography (micro-CT) to confirm the fiber locations at the end of the experiment^56^ (Fig. 2c, Extended Data Fig. 2).

**Fig. 2.**
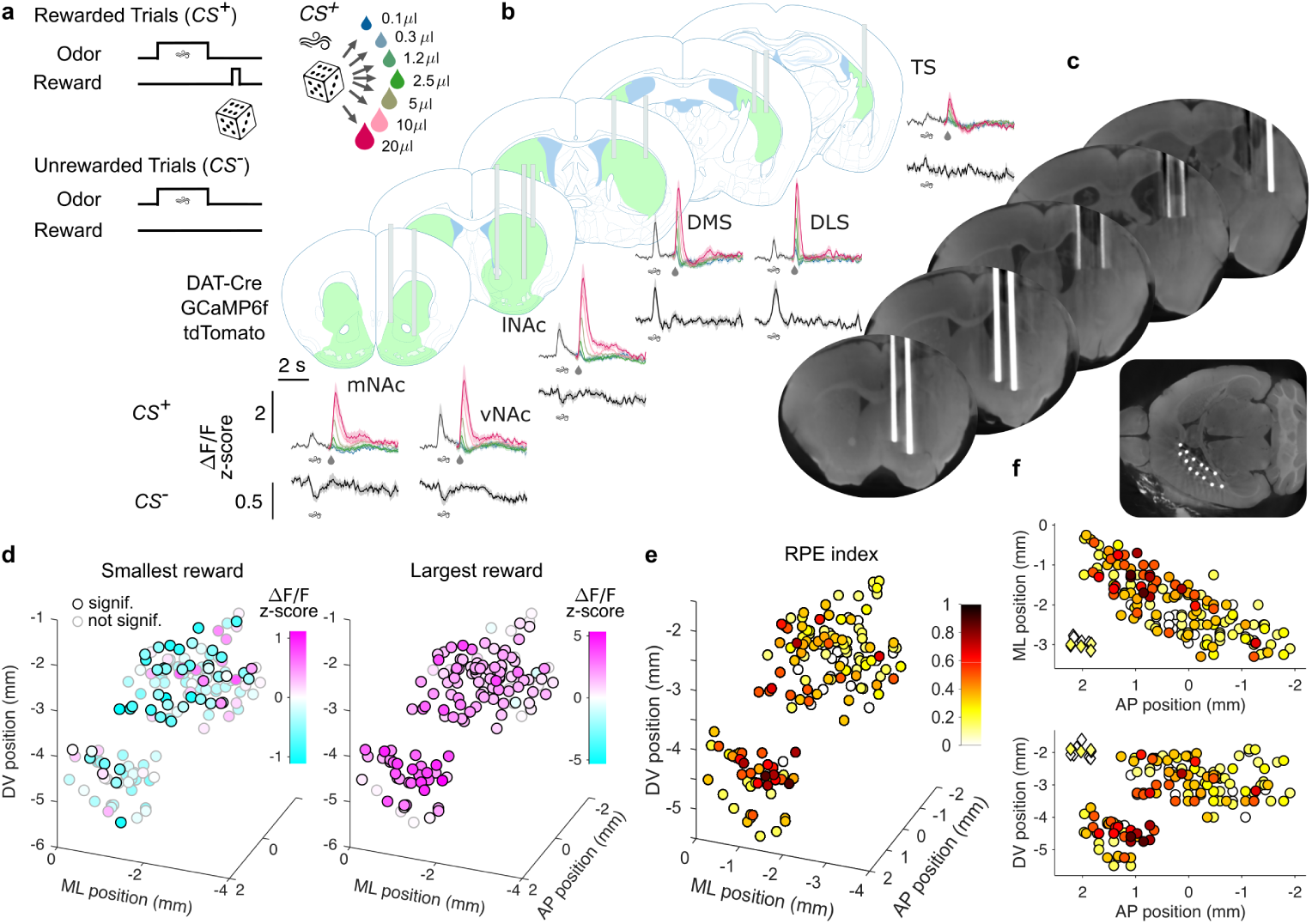
Anatomical organization of canonical RPE signals across the striatum. **a**, Classical conditioning task used to characterize the activity of positive and negative prediction errors across the striatum. **b**, Simultaneous multi-fiber recording of calcium activity in dopamine axons across the striatum with example average fluorescence traces for *CS*^+^ (top) and *CS^−^* (bottom) trials in 6 regions (shaded areas represent s.e.m across mice: *n* = 11). **c**, Micro-CT scan of an example mouse showing fiber placement in target regions aimed at in **b**. **d**, Average fluorescence response (z-scored Δ*F/F*_0_) to the smallest (left) and largest (right) reward measured in each fiber (14 fibers in each of *n* = 11 mice) matched to its recording location in the striatum in relation to Bregma: AP anterior-posterior axis, ML medial-lateral axis, DV dorso-ventral axis. Fibers with black edge show significant modulation from baseline (*P <* 0.05 Wilcoxon sign rank test), fibers with grey edge signal non-significant modulation from baseline. **e**, RPE index mapped to fiber location in the striatum (all edges are shown in black; only significant criteria contribute to an increase in RPE index, see Methods). **f**, Same data as in **e** shown in the AP-ML projection (top, horizonal view) and in the AP-DV projection (bottom, sagittal view). Diamonds around the (2*AP,* 3*ML,* 2*DV*) coordinates correspond the control ROIs in the cameras’ fields of view, not to real fibers in the brain.

Mice were trained in the variable reward task^2,60^ in which one odor (*CS*^+^) predicted a drop of water delivered after a fixed delay, with its volume drawn on each trial from a predefined distribution (Fig. 2a). A second odor predicted no reward (*CS^−^*). At all recording sites, dopamine axon responses increased with reward size, although they were generally small and variable in the TS (Fig. 2b, Extended Data Fig. 3). Across the entire striatum, the smallest reward tended to elicit a negative response while the largest reward elicited a positive response, a pattern that became less salient towards the posterior striatum (Fig. 2d, Extended Data Fig. 3).

**Fig. 3.**
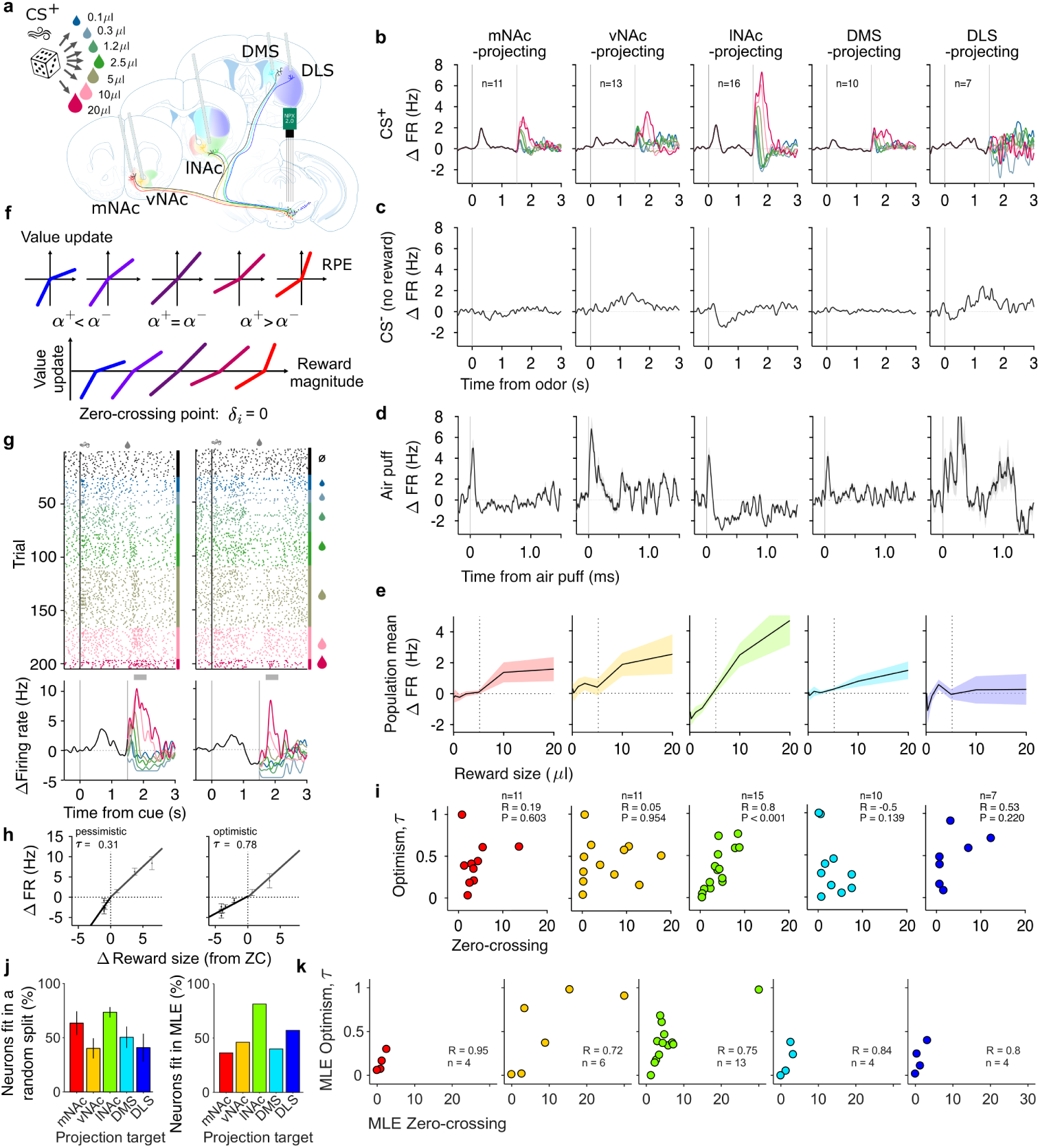
lNAc-projecting dopamine neurons are compatible with distributional RL. **a**, Experimental approach for recording of projection-identified dopamine neurons using antidromic optotagging from 5 striatal locations: mNAc (*n* = 11), vNAc (*n* = 13), lNAc (*n* = 16), DMS (*n* = 10), DLS (*n* = 7). **b**, Projection-specific population average activity during *CS*^+^ trials. Traces after outcome delivery are color-coded according to reward size. **c**, Projection-specific population average activity during *CS^−^* trials. **d**, Projection-specific population average activity during unpredicted air puff delivery (shaded areas represent s.e.m across neurons). **e**, Projection-specific population response function to reward size during *CS*^+^ trials (vertical dotted line signals mean of the distribution, i.e. expected value). **f**, Schematic of diverse response functions for dopamine neurons in the distributional RL framework: piece-wise linear functions with variable slopes (top) lead to asymmetric value updates causing diversity in zero-crossing points (bottom). **g**, Raster plot (top) and PSTH (bottom) of two example neurons projecting to the lNAc recorded in the same session. Colors match reward size in *CS*^+^ trials; *CS^−^* trials are shown in black in the raster plot but not shown in the PSTH. In the PSTH, black trace shows the average response across all *CS*^+^ trials before reward delivery, and grey bar signals time interval used for analysis of zero-crossing and optimism. **h**, Empirical estimation of zero-crossing point and optimism for the two example neurons in **g**: baseline-subtracted activity for each reward size is plotted against the difference between the delivered reward size and the estimated zero-crossing point. Lines show linear fit to the data for positive and negative RPEs (slopes are used to calculate optimism level). **i**, Relationship between empirical zero-crossing point and optimism for each projection-defined population of dopamine neurons, with corresponding Pearson’s correlation coefficient. **j**, Percentage of neurons for which we can fit zero-crossing and optimism parameters using the empirical method (left) and using a more stringent model-based approach to simultaneously estimate zero-crossing point and optimism through a maximum likelihood fitting procedure (right, see Methods). **k**, Relationship between model-inferred zero-crossing point and optimism for each projection-defined population of fitted dopamine neurons.

Cue responses were significantly larger to *CS*^+^ than to *CS^−^* across all striatal regions except DLS (Extended Data Figs. 3, 4). Notably, negative responses to *CS^−^* were only found in the ventral striatum and anterior dorsal striatum.

To systematically characterize the anatomical organization of canonical RPE signaling, we quantified eight complementary features of dopamine responses across three behavioral tasks. We then combined these features into an RPE index to compare the prevalence of canonical RPE signaling across striatal territories. These features were: 1) a decrease of reward response during learning (Extended Data Figs. 5, 6), 2) an increase of cue response during learning (Extended Data Figs. 5, 6), 3) a temporal shift of response from the reward time to the cue during learning (Extended Data Figs. 5, 6), 4) a negative correlation between cue response and delay to reward in a cued delay task^61^ (Extended Data Fig. 7), 5) negative response to a reward smaller than the expected value of the reward distribution (Fig. 2d, Extended Data Fig. 4), 6) a negative response to *CS^−^* (Extended Data Fig. 4), 7) a negative response to an air puff (Extended Data Fig. 8), 8) a negative response to a cue that predicts an air puff (Extended Data Fig. 9). We found that the RPE index was highest in the nucleus accumbens, particularly in the lNAc, and there was a significant anatomical gradient in the anterior-posterior, medio-lateral, and dorso-ventral anatomical axes (Extended Data Fig. 10). These results confirm the predominance of canonical RPE signals in the ventral striatum, especially in the lateral nucleus accumbens^55,62,63^, while demonstrating that the full complement of canonical RPE features becomes progressively less prevalent towards the dorsolateral and posterior striatum^51,64,65^.

**Fig. 4.**
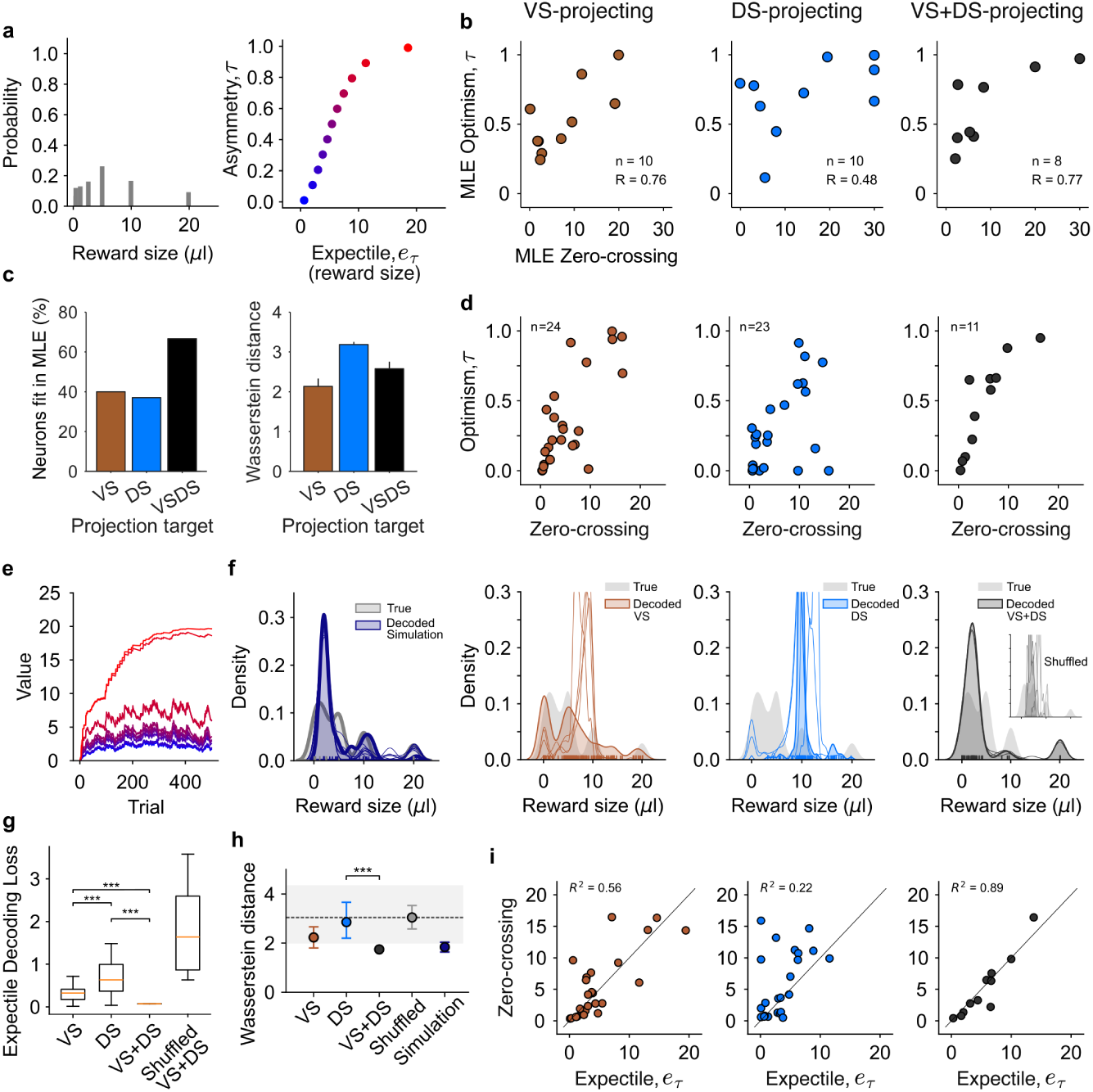
Reward distribution information reaches the ventral and dorsal striatum in a single mouse. **a**, Theoretical probability distribution for rewards delivered following *CS*^+^ odor (left) and corresponding inverse expectile function using 11 expectiles (right). **b**, Relationship between model-inferred zero-crossing point and optimism for each projection-defined population of fitted dopamine neurons (left: tagged only by VS fiber(s) in brown, *n* = 10; middle: tagged only by the DS fiber in blue *n* = 10; right: tagged by both VS and DS fibers in black, *n* = 8). **c**, Percentage of neurons that do fit the model (left), and Wasserstein distance between decoded and true distributions (mean x00B1± SD over 10 decoding runs, right), for each projection target. **d**, Empirically-estimated optimism level versus zero-crossing point for each population of projection-defined dopamine neurons (VS-projecting: *n* = 24, positive correlation *P <* 0.001; DS-projecting: *n* = 23, positive correlation *P* = 0.01; VS+DS- projecting: *n* = 11, positive correlation with *P <* 0.001). **e**, Example Rescorla-Wagner simulation of value-learning of this reward distribution using 11 value predictors. **f**, Expectile decoding of the reward distribution shape from 10 simulations of 11 value predictors after 500 trials of learning (right, each line is one simulation), from five example subsets of 11 randomly sampled VS-projecting neurons (brown, each line is one draw), from five example subsets of 11 randomly sampled DS-projecting neurons (blue, each line is one draw) and from the 11 VS+DS-projecting neurons (black, each line is one decoding run, inset in grey: decoding from VS+DS shuffled values). Shaded distributions correspond to the decoding run with the smallest loss, gray shade in the background represents the true distribution density from **a**. **g**, Expectile decoding loss per projection target. For both VS and DS, decoding was performed in 1000 random subsets of 11 neurons from these populations. For VS+DS neurons, 10 decoding runs were performed, and 10 shuffles of the optimism and zero-crossing values were used for decoding of shuffled control. Decoding loss differed across VS, DS, and VS+DS populations (overall permutation test, *P <* 0.001). Post hoc pairwise permutation tests with Holm correction showed that all populations differed from one another *P <* 0.001. **h**, Wasserstein distance between the decoded and true distributions. Dots represent means, error bars indicate 95% bootstrap confidence intervals for the experimental populations and the 2.5*^th^*–97.5*^th^* percentile interval for simulations, gray shaded band indicates the 2.5*^th^*–97.5*^th^*percentile interval of the shuffled VS+DS control, and the dashed line indicates its mean. Permutation tests were used to compare VS, DS and VS+DS populations (overall permutation test *P* = 0.0128). Post hoc pairwise permutation tests with Holm correction showed that DS differed from VS+DS (*P <* 0.001), but not the other two pairs. **i**, Coefficient of determination, *R*^2^, between the zerocrossing point and the predicted expectile, *e_τ_*, for the optimism level of each neuron.

Together, these experiments provide a comprehensive, spatially resolved map of dopamine signaling across the striatum, systematically characterizing eight features of canonical RPE activity across multiple behavioral tasks (Fig. 2, Extended Data Figs. 3–10). This mapping reveals a graded anatomical organization of RPE signaling, with the lNAc emerging as the region exhibiting the most complete repertoire of canonical RPE features, including robust negative responses to worse-than-expected outcomes. Importantly, the partial expression of these features in other striatal territories highlights the functional heterogeneity of dopamine signaling and suggests that canonical RPEs coexist with other types of dopamine signals across the striatum.

### 2.3 lNAc-projecting dopamine neurons are compatible with distributional RL

Fiber photometry cannot distinguish whether dopamine signals recorded within a striatal region arise from a homogeneous population of axons or from intermixed axons of functionally distinct dopamine populations. We therefore performed electrophysiological recordings of projection-defined dopamine neurons at the single neuron level, combining high-density Neuropixels recordings of somatic spiking with optogenetic identification through antidromic activation of their axons^34^ (Extended Data Fig. 11).

We adapted our custom implant to carry 5 optical fibers for antidromic excitation in the mNAc, vNAc, lNAc, rDMS and iDLS (Extended Data Fig. 11a), and trained mice in the variable reward task (Fig. 3a). The average response of all projection-defined populations, except the DLS-projecting one, showed graded responses to reward size (Fig. 3b,e). However, only the lNAc-projecting population showed negative responses to rewards smaller than the expected value of the reward distribution (5.4*µl*, Fig. 3b,e), and to the *CS^−^* (Fig. 3c). The lNAc-projecting population thus carries balanced, bidirectional RPEs, consistent with both traditional and distributional RL, whereas the DLS-projecting population does not fit the RL framework for reward-based value learning. The mNAc-, vNAc-, and DMS-projecting populations are ambiguous: the lack of negative responses to the small rewards could reflect an inability to signal negative prediction errors, or extreme pessimism in distributional RL, such that most reward deliveries elicit positive RPEs. To distinguish these possibilities, we delivered unpredicted air puffs, which should elicit negative responses in any population signaling negative RPEs. Unpredicted air puff caused a brief, short-latency, excitation in all populations, consistent with a salience-like response reported previously^66,67^, but the delayed component encoding value^2,68,69^ was negative only in the lNAc-projecting population (Fig. 3d). Consistent with the photometry results, the lNAc-projecting dopamine population is therefore the main carrier of bi-directional RPEs, and the strongest candidate for implementing distributional RL in the striatum.

We next fit the distributional RL model to each dopamine neuron population and estimated the key model parameters (zero-crossings and optimism) using previously developed methods^2,36^. First, we estimated the zero-crossing point (i.e. the reward magnitude that elicits zero RPE) using the response function of each neuron. Second, we obtained its optimism, *τ*, from the slopes of positive and negative responses (equivalent to *α*^+^ and *α^−^*, respectively, Fig. 3f, g, h). Distributional RL predicts that these two variables are positively correlated^2,23^(Fig. 1b).

We estimated the correlation between these two variables in 50 random partitions of the data, and observed that only the lNAc-projecting population showed positive and significant correlations (Extended Data Fig. 13d). When we average the zero-crossing and the optimism level *τ* across these splits to obtain their Pearson’s correlation, only the lNAc-projecting population shows a significant positive correlation between these two variables. Other populations did not show this key signature of distributional RL (Fig. 3i). Thus, our analysis suggests that the lNAc-projecting population is the ideal candidate for the implementation of distributional RL in the dopamine system.

We confirmed these results in a separate analysis in which all dopamine neurons are pooled before any fitting takes place, instead of separating them by projection target before the analysis (Extended Data Fig. 14).

Our analysis so far relied on the estimation procedure introduced in our previous study^2^, in which the zero-crossing point is first estimated from the neural responses and the positive and negative slopes are subsequently fit on either side of this point. To avoid circularity, zero-crossing points and slopes are estimated from independent halves of the data before computing their correlation. Although statistically unbiased, this approach uses only half of the available trials for each estimate, reducing statistical power, particularly for infrequently delivered reward sizes.

To overcome this limitation, we developed a maximum likelihood estimation (MLE) framework that jointly estimates zero-crossing and optimism from all available trials (see Methods). This method finds the parameters that fit the entire dataset best, increasing robustness for low-data regimes. Because neither parameter is conditioned on the other, any correlation between them cannot be an artifact of the fitting procedure. We therefore use it as a normative companion for the results from the standard fitting procedure. This method also provides a more stringent criterion for whether a neuron has a zero-crossing point corresponding to an expectile of the distribution: a neuron is considered fitted only if its response increases with reward on both sides of its zero crossing point. Our MLE procedure fitted the activity of most lNAc-projecting dopamine neurons (81.3%), but fewer neurons projecting to mNAc (36.4%), nNAc (46.2%), DMS (40%), and DLS (57.1%) (Fig. 3j). A substantial fraction of neurons projecting outside of lNAc appeared to lack the bi-directional, graded responses required by distributional RL (Fig. 3e, Extended Data Fig. 12b). The fitted neurons in these populations nevertheless showed a strong correlation between zero-crossing and optimism (Fig. 3k), suggesting that they may also contribute to the reward distribution representation.

Together, these results identify the lNAc as a hotspot for receiving bi-directional RPE signaling and for the implementation of distributional RL.

### 2.4 Broadly-projecting mesolimbic dopamine neurons broadcast outcome distribution information

Our results so far indicate that the response of lNAc-projecting dopamine neurons is the most consistent with distributional RL. However, distributional signatures have been observed in the activity of striatal neurons more broadly in the anterior ventral and dorsal striatal regions^3^. Notably, a few of our dopamine neurons were antidromically idendified from more than one striatal region, most commonly including lNAc (Extended Data Fig. 11e). This is consistent with previous reports of axonal collateralization by lNAc-projecting dopamine neurons^70^ and recent anatomical identification of sparse but broadly projecting dopamine populations^71^. We therefore asked whether these broadly projecting neurons also encode reward distribution information.

We implanted three larger optical fibers in the striatum, two targeting the lNAc and one the dorsal striatum (DS), and increased photostimulation power for more efficient optotagging (Extended Data Fig. 11f,i). In one mouse we identified 64 neurons (Extended Data Fig. 16a,b), despite the fact that the optical fibers aimed at lNAc reached neighboring locations around our targets, namely the lateral core of the NAc and the ventral striatum. We will therefore label these fibers as VS for lateral ventral striatum (Extended Data Fig. 16c).

Out of the 64 identified dopamine neurons, 12 were optotagged by one or both VS fibers as well as by the DS fiber (we will refer to this population as “VS+DS population”). The remaining neurons were tagged only by (one or two) VS fiber(s) (“VS-projecting population”), or only by the DS fiber (“DS-projecting population”). The VS+DS population can provide shared information across the ventral and dorsal striatum (Extended Data Fig. 17). Applying the maximum likelihood fitting procedure (Fig. 4b) we found that the VS+DS population had the largest fraction of neurons meeting requirements of expectile coding: this population had a significant correlation between zero-crossing points and optimisms, comparable to that fitted in the VS-projecting population (Fig. 4c). An expectile decoder^2^ recovered the reward distribution least successfully from the DS-projecting population (Extended Data Fig. 18a,e). Furthermore, bootstrapped correlation between zero-crossing and optimism was highest in the VSand VS+DS-projecting populations (Extended Data Fig. 18d). Finally, we also applied the same empirical method as before (in Fig. 3)^2^ to estimate zero-crossing and optimism from these data, and obtained qualitatively similar results to those with the MLE method (Fig. 4b,d). This empirical calculation resulted in the estimation of zero-crossing and optimism for 11 neurons from the VS+DS population (Fig. 4d). To test whether 11 value predictors are enough to represent the reward distribution, we used the equation in Fig. 1d to run multiple simulations of distributional learning in groups of 11 value predictors (Fig. 4e). We then run the expectile decoder in these simulated populations of 11 value predictors to obtain the shape of the decoded distribution in such a data limited regime (Fig. 4f). Running the decoder in the VS+DS population shows that this small population contains similar information about the reward distribution shape as the simulation of 11 value predictors. To evaluate the decoding capacity of the VS-only and DS-only-projecting populations, we performed decoding in 1000 random draws of 11 neurons from these populations (out of 24 and 23 neurons from the VS-projecting and DS-projecting populations, respectively). We found that, depending on the draw, groups of VS-projecting dopamine neurons contain detailed information about reward distribution, or have little information about it. By contrast, no draw from the DS-projecting population captures the distribution shape information (Fig. 4f, 5 decoded shapes are included in the figure as examples). We quantified the quality of the decoding using the expectile decoding loss and the Wasserstein distance between the decoded and true distributions. These were smallest for the VS+DS population and largest for the DS-projecting population (Fig. 4g,h). Finally, for each neuron, we matched its zero-crossing point to the theoretical expectile at the same optimism level (Fig. 4i). The highest coefficient of determination was observed in the VS+DS population, suggesting that this population specifically shares reward distribution shape information between the ventral and dorsal striatum.

Our findings reveal that, besides lateral VS-projecting dopamine neurons, a population of dopamine neurons projecting to both VS and DS conveys information about the shape of the reward distribution, broadcasting distributional reward prediction errors to both the ventral and dorsal striatum.

### 2.5 Distribution-encoding mesolimbic dopamine neurons neurons form an anatomical gradient of optimism in the VTA

Both lateral VS-projecting (Figs. 3, 4) and VS+DS-projecting (Fig. 4) dopamine populations exhibit structured heterogeneity consistent with distributional RL, unlike the DS-projecting population. Both populations span the full spectrum of optimism, from pessimistic to optimistic prediction errors, raising an important question: how are these distinct RPE signals kept separate to selectively update their corresponding striatal value predictions (Fig. 1)? Previous work suggests that D1- and D2-expressing striatal neurons preferentially learn from positive and negative prediction errors, respectively^3^, providing a potential mechanism for separating optimistic and pessimistic updates in the striatum. How such functional separation is maintained in the midbrain, however, remains unclear. Dabney et al. (2020) proposed that distributional learning could be preserved through topographically organized recurrent connectivity, whereby striatal value predictors preferentially connect to dopamine neurons in the same regions from which they receive RPEs^2^. Motivated by this prediction, we examined the anatomical organization of optimism among dopamine neurons.

We reconstructed the anatomical locations of the recorded dopamine neurons (Fig. 5a) and found that VS- and VS+DS-projecting populations were located in the ventral tegmental area (VTA) and medial substantia nigra pars compacta (SNc). Univariate analysis revealed a mediolateral gradient of optimism, with optimistic neurons located more medially and pessimistic neurons more laterally (Fig. 5b). To determine whether this relationship was independent of the other anatomical coordinates, we performed multiple linear regression, which revealed a significant association between optimism and three-dimensional anatomical location (*R*^2^ = 0.396, *F*_3,30_ = 6.56, *P* = 0.0015, *n* = 34 neurons from one mouse). Optimism decreased along the mediolateral axis (*β*_ML_ = *−*0.631, *P* = 0.0065) and increased along the dorsoventral axis (*β*_DV_ = 0.936, *P* = 0.0158), after controlling for the other coordinates, whereas no significant independent association was observed along the anteroposterior axis (*β*_AP_ = 0.230, *P* = 0.475; Extended Data Fig. 21). Dopamine neurons located in the lateral SNc, by contrast, projected exclusively to the dorsal striatum (Extended Data Fig. 22).

**Fig. 5.**
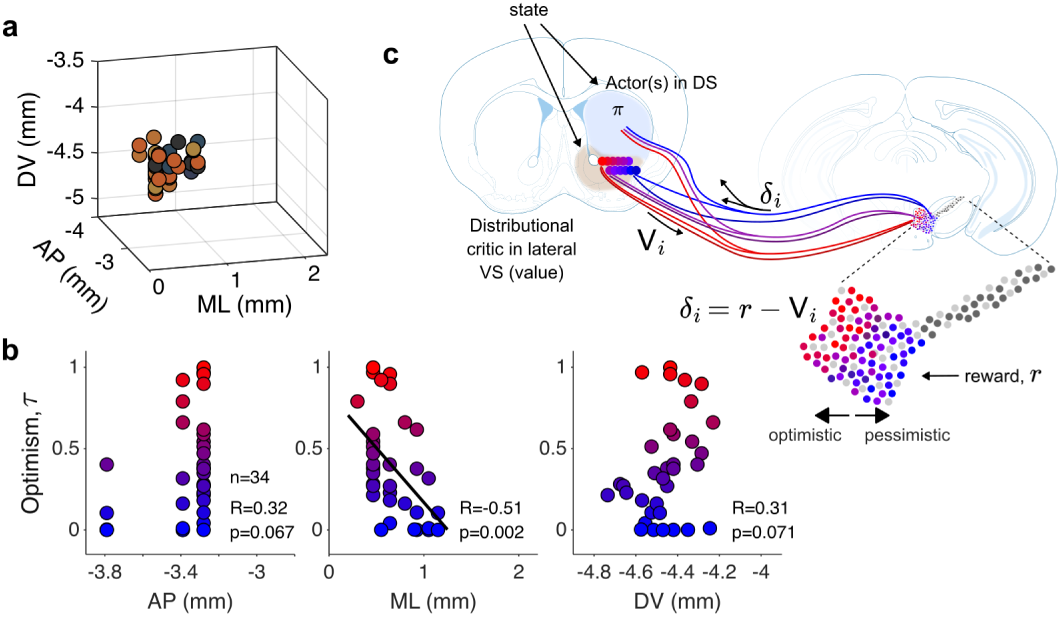
Anatomical gradient of optimism in the VTA / medial SNc. **a,** Anatomical location of VS and VS+DS-projecting dopamine neurons analysed in Fig. 4. Brown circles represent neurons projecting only to VS, dark grey and black circles represent neurons projecting to VS+DS (color is set based on the dot product of the optotagging information with the color representing each fiber). **b**, Optimism level of the neurons shown in **a** as a function of anatomical axis: anterior-posterior (left), medio-lateral (center), dorso-ventral (right). **c**, Schematic summarizing the neural circuit organization of dopamine-dependent distributional RL: dopamine neurons are organized in a gradient of optimism in the VTA and loop with striatal neurons in the lateral VS to enable learning of the outcome distribution, while also sending axonal collaterals to DS, broadcasting this information. This configuration is akin to a distributional actor-critic model of the basal ganglia.

Together, these findings identify an anatomically organized population of mesolimbic dopamine neurons capable of broadcasting distributional RPEs to both ventral and dorsal striatal circuits, providing a potential substrate for coordinating distributional value learning and behavioral policy updates.

## 3 Discussion

Here, we sought to uncover the circuit-level implementation of distributional RL by dopamine neurons (Fig. 1). Canonical bidirectional RPE signaling is necessary but not sufficient for distributional RL: zero-crossing points and optimism must vary across neurons and be positively correlated^2^. We therefore first mapped the prevalence of canonical RPE signals across the striatum using fiber photometry, followed by projection-specific single-unit recordings to identify signatures of distributional RL. We found that canonical bidirectional RPE signals are most prevalent in the lateral nucleus accumbens (lNAc; Fig. 2), and that dopamine neurons projecting to this region exhibit the population-level signatures required for distributional RL (Fig. 3). Although other striatal regions also receive signals exhibiting some canonical RPE features, distributional signatures were weaker or absent in neurons projecting exclusively to these regions. These results are consistent with our previous finding that distributional value representations in striatal neurons are strongest in the lNAc but also detectable elsewhere^3^. Importantly, we identified a population of dopamine neurons projecting to both the lateral ventral striatum, including the lNAc, and the dorsal striatum, providing an anatomical substrate for the broader distribution of these signals. These broadly projecting neurons exhibit structured heterogeneity that enables the reconstruction of the reward distribution. Notably, this representation does not arise solely from pooling neurons across animals, as reward distribution information could be decoded from neurons recorded in a single mouse (Fig. 4).

Together, these findings reveal a previously uncharacterized functional organization of broadly projecting dopamine neurons^70,72,73^, providing a circuit architecture through which distributional prediction errors can be shared across ventral and dorsal striatal regions.

Importantly, both lateral VS-projecting and VS+DS-projecting dopamine populations span the full spectrum of optimism, from pessimistic to optimistic prediction errors, raising the question of how these distinct teaching signals remain functionally segregated. We found that optimism is organized along a mediolateral gradient, with an additional dorsoventral component, across the VTA and medial SNc (Fig. 5), providing a potential anatomical substrate for maintaining distinct inputs from optimistic and pessimistic value predictors. The mechanisms underlying this organization remain unknown: such gradient in optimism does not appear to be explained by baseline firing rates alone^2,74^ (Extended Data Fig. 13b), and the lateral habenula–rostromedial tegmental nucleus pathway, which conveys inverse RPEs and inhibits dopamine neurons^75–77^, does not readily account for the full spectrum of optimism^74^. One possibility is that spatially biased inputs differentially shape dopamine responses to positive and negative outcomes. Such organization could arise from distinct feedback pathways involving D1 and D2 striatal projection neurons and the ventral pallidum: our previous work suggests that asymmetric dopamine-dependent plasticity of D1 and D2 striatal neurons could transform heterogeneous dopamine prediction errors into distinct distributional value predictions^3^, while the anatomical organization of dopamine neurons in the midbrain may help preserve the correspondence between these predictions and their teaching signals. The role of other inputs exhibiting mediolateral organization, including serotonergic projections from the dorsal raphe^78^, remain to be tested.

Identifying the circuit organization of distributional RL presents substantial experimental and analytical challenges. Projection-specific recordings require sampling hundreds of midbrain neurons to identify the small fraction (4–6%) that can be reliably antidromically tagged, limiting the number of neurons available for analysis. Furthermore, estimating zero-crossing points and optimism from noisy single-neuron responses and limited trial numbers is challenging, particularly when responses to negative prediction errors are weak. Different fitting approaches can yield differences in absolute parameter estimates and in the fraction of neurons meeting the criteria for distributional coding. Nevertheless, the convergence of population-level results across complementary fitting approaches supports the robustness of our conclusions about the anatomical organization of distributional RPE signals.

Our findings have three broader implications. First, the anatomical gradient of optimism provides a potential substrate for maintaining distinct distributional learning signals, which could support risk-sensitive learning and decision-making^41^. Second, the broad axonal projections of distribution-encoding dopamine neurons suggest that the DS receives functionally distinct dopamine signals from different midbrain sources: distributional RPEs from VTA and medial SNc neurons, alongside other dopamine signals originating in lateral SNc^64^. Third, the broadcasting of distributional RPEs to DS raises the possibility that these signals contribute not only to value learning in VS but also to the learning and updating of behavioral policies in DS.

Indeed, dopamine RPEs have long been proposed to provide teaching signals for both value learning and reward-optimizing actions, i.e., behavioral policy^30^. Actor–critic models of RL provide one such formalization, inspiring a view of the basal ganglia in which VS is proposed to function as a *critic*, learning state values, while DS acts as an *actor*, learning behavioral policies^7,8,79^. Our findings suggest a distributional extension of this architecture (Fig. 5c): lateral VS circuits may implement a distributional critic, while broadly projecting dopamine neurons broadcast distributional prediction errors to DS circuits involved in action learning. These signals could reinforce action representations and shape subsequent choices^35,80–83^. Determining whether and how heterogeneous distributional RPEs contribute to policy learning will be an important direction for future research.

Distributional critics have already been incorporated into artificial RL agents performing complex continuous-control tasks, including humanoid locomotion and robotic manipulation^22,24^, as well as hierarchical tasks such as autonomous racing^25^. An important next step will be to determine whether broadcasting heterogeneous distributional prediction errors to action-learning circuits enables behavioral policies to adapt more efficiently to changes in environmental statistics, internal states, and behavioral goals.

## 5 Methods

### 5.1 Experimental procedures

#### Mice

A total of 25 adult C57BL/6J mice (Jackson Laboratory, 000664) of both sexes were used in this study. All animals were DAT-Cre mice (*Slc6a3 ^tm^*^1.1(*cre*)*Bkmn*^, Jackson Laboratory, 006660)^84^. These mice could be single transgenic, or double transgenic by crossing DAT-Cre with _Ai32 (B6;129S-*Gt(ROSA)*26_*_Sortm_*_32(_*_CAG−COP_* _4_*_∗H_*_134_*_R/EY FP_* _)_*_Hze_*_/J, Jackson_ Laboratory, 012569)^85^, or double transgenic after crossing with Ai148D (B6.Cg_Igs7_*_tm_*_148.1(_*_tetO−GCaMP_* _6_*_f,CAG−tTA_*_2)_*_Hze_*_/J, Jackson Laboratory, 030328)58, or_ triple transgenic DAT-Cre::Ai148::tdTomato by crossing further with Ai14 (B6.Cg*Gt(ROSA)26Sor^tm^*^14(*CAG*^*^−tdT^ ^omato^*^)*Hze*^*/* J, Jackson Laboratory, 007914)^59^. Mice were aged two to four months at the time of surgery (the first experimental procedure). Animals were housed on a 12 hr dark/12 hr light cycle and performed the tasks during the dark period. Ambient temperature was kept at 755*F*, and humidity was kept below 50%. Animals were group-housed (2–5 animals/cage) until surgery, then individually housed throughout training and testing. All procedures were performed in accordance with the National Institutes of Health Guide for the Care and Use of Laboratory Animals and approved by the Harvard Institutional Animal Care and Use Committee (IACUC).

#### Surgeries

All surgical procedures were performed under aseptic conditions. Mice were anesthetized with isoflurane (4% for induction, 1–2% for maintenance at 0.5–1 L.min*^−^*^1^), and analgesia was given pre-operatively using buprenorphine (0.1 mg.kg*^−^*^1^, intraperitoneal (i.p.)). Following head-fixation in the stereotactic frame, hair removal, and disinfection of the scalp with 70% ethanol and 10% povidone-iodine, a local anesthetic (lidocaine (2%)/bupivacaine (0.5%) 1:1 mixture, subcutaneous (s.c.)) was applied at the incision site. The skull was leveled in the anterior-posterior axis, such that bregma and lambda were at the same dorso-ventral position. The coordinates for the craniotomies (for viral injection and/or for implant introduction) were marked on the skull using a fine-tipped pen or the tip of the dental drill. The coordinates used were the following:

**Table 1.**
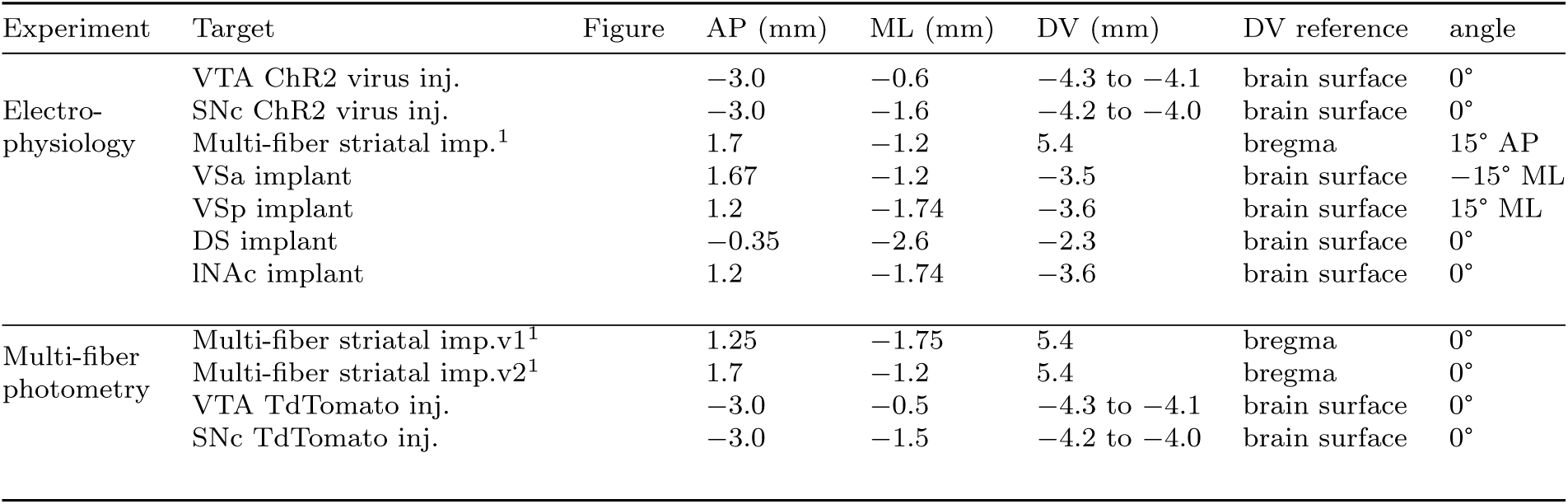
Stereotactic coordinates used. ^1^Given the fixed position of of the fibers in the multi-fiber implant, only one fiber was used as reference for alignment and implantation

A custom-made titanium head plate was placed on the well-cleaned and dried skull with adhesive cement (C&B Metabond, Parkell), containing a small amount of charcoal powder. Following head-late placing, the craniotomies were performed with a high-speed dental drill.

When viral injections were performed, the viral solution was injected at a rate of approximately 20 nl.min*^−^*^1^ using glass pipettes (Drummond, pulled on a heating puller: PC-10, Narishige) and a manual plunger microinjector (MO-10, Narishige). Typically, the viral solution was dispersed within a 200*µm* line below and above the target DV coordinate. The pipette was withdrawn at least 10 minutes after the end of injection.

For electrophysiological recordings with Neuropixels probes, optical fibers were inserted in the striatum. In the case of the 5-fiber custom implant (custom B283-2005 HDFC 200/250-0.66 HD19 0005, Doric Lenses, Extended Data Fig. 11a), a single craniotomy was made over the left striatum and the implant was lowered slowly inside the moist brain. This implant contained 5 optical fibers with a core of 200*µm* and NA 0.66. In the case of the 3-fiber and 1-fiber implantations in the striatum (Fig. 11c,d), we used single 400*µm* core optical fibers with NA 0.48 (MFC 400/430-0.48 5mm MF1.25 FLT for VS targets and MFC 400/430-0.48 3mm MF1.25 FLT for DS, Doric Lenses). After fiber implantation, a ground pin coated with a lead-tin alloy (Mill-Max) was implanted above the contralateral side of the skull and cortex. Finally, a 3D printed plastic well was glued on top of the headplate with 5-minute epoxy.

For multi-fiber photometry recordings, a custom 14-optical-fiber cannula implant (HDFC 200/250-0.66 HD19 0002 manufactured by Doric Lense) was inserted above the striatum using a custom holder (SCH 2.5 and FCA 2.5 HD19, Doric Lenses), after drilling a single, large, bean shaped craniotomy, whose most anterior and posterior coordinates were, respectively, (2.0*mm* AP,*−*0.5*mm* ML) and (*−*1.5*mm* AP,*−*3.75*mm* ML). The implants were secured with C&B Metabond adhesive cement (Parkell) and dental acrylic (Lang Dental).

Following surgery, mice were given saline (0.5*ml*) and analgesia (ketoprofen, 20*mg/kg*, injected subcutaneously), placed inside a clean cage and recovered over a heating pad. Mice were kept single housed and given analgesia twice daily for at least 48 hours. When viruses were injected, we waited at least 3 weeks to start the experiments.

#### Viruses

For electrophysiological experiments in which single transgenic DAT-Cre mice were used (*n* = 10, Fig. 11c,d), a virus containing ChR2 was injected in the VTA and SNc: rAAV5-EF1a-DIO-hChR2(H134R)-eYFP-WPRE-pA (titer: 5.3 *×* 10^12^ gc/ml, University North Carolina (UNC) Vector Core). In multi-fiber photometry experiments in which double transgenic mice (DAT-Cre::Ai148, *n* = 3) were used (instead of triple transgenic DAT-Cre::Ai148::tdTomato), tdTomato was injected in the VTA and SNc: AAV5-CAG-Flex-tdTomato (titer: 4.8 *×* 10^12^ gc/ml, UNC Vector Core). In two-photon calcium imaging experiments in which DAT-Cre mice were used (*n* = 2, instead of DAT-Cre::Ai148), we injected GCaMP7f in the VTA and SNc: AAV5-CAG-FLEX-jGCaMP7f (300*nl*, titer: 1.8 × 10^13^ particles per ml, UNC Vector Core, custom preparation)^86^.

#### Head-fixed behavioral tasks

Across all neural recording modalities, mice were water-deprived in their home cage 1–2 days before the start of behavioral training, two or more weeks after surgery. During water deprivation, each mouse’s weight was maintained above 85% of its original value.

Mice were habituated to the head-fixed setup by receiving water every 4*s* (6 *µ*l drops) for 3 days, after which association between odors and outcomes started. A mouse lickometer (1020, Sanworks) was used to measure licking as infrared beam breaks.

Water valves (LHDA1233115H, The Lee Company) were calibrated, and a custommade olfactometer based on a valve driver module (1015, Sanworks) and a valve mount manifold (LFMX0510528B with LHDA1221111H valves, The Lee Company) was used for odor delivery. All components were controlled through a Bpod state machine (1027, Sanworks).

Odors were diluted in mineral oil (Sigma-Aldrich) at 1 : 10, and 30*µ*l of each diluted odor was placed inside a syringe filter (2.7*µ*m pore size, 6823-1327, GE Healthcare). Odorized air was further diluted at 1 : 10 and delivered at 1,000*ml.min^−^*^1^.

Odors used for association in the rewarded or in the air puff tasks were randomly assigned from the following list of odors: isoamyl acetate, p-cymene, ethyl butyrate, (+)-carvone, (*±*)-citronellal, *α*-ionone, L-fenchone, caproic acid, eugenol, 1-hexanol, ethyl acetate, octanoic acid, 1,4-cineole, hexyl tiglate. For the rewarded odor, a reward was delivered after a trace period of 0.5*s* in electrophysiology experiments, and of 0.5*s* and/or 2*s* in multi-site fiber photometry experiments. In the variable reward magnitude task, one of these odors was associated with a distribution of reward sizes (0.1, 0.2, 1.25, 2.5, 5.0, 10.0, 20.0 *µ*l) while a second odor was not paired with any outcome (nothing). The water delivery of fixed reward size was 5.4*µl* (Extended Data Fig. 5 and 6) or 4*µl* (Extended Data Fig. 7). In the cued delay task, 4 odors predicted different delays to a fixed reward deterministic delivery. The delays used were 1.25*s,* 2.5*s,* 4.0*s,* 6.5*s*. In this task (as well as in the air puff association task), a blue LED preceded the odor delivery by 1.5*s* to eliminate potential surprise components in the cue response^61^. For the air puff delivery we used a pressure regulator (LRP-1/4-4, Festo) and a solenoid valve (MHE2-MS1H-3/2G-M7-K, Festo) controlled by Bpod. Air puffs had a duration of 30*ms* and a pressure of 20*psi*. In the air puff association task, air puff was delivered 1.25*s* after odor onset. Mice completed one head-fixed behavioral session per day.

#### Acquisition of multi-fiber photometry signals

To collect fluorescence signals from dopamine axons across the striatum we used a bundle-imaging fiber photometry system with two integrated CMOS cameras and 3 integrated light-emitting diodes (LEDs) (BFPD, USB H4, PR BFP, LEDD 4, and BFMC6 LED(410-420) LED(460-490) CAM(500-550 LED(555-570) CAM(580-680), Doric Lenses). The three LEDs were triggered sequentially and illuminated all of the fibers implanted in the brain through a bundle patchcord (HDP(19) 200/245/LWMJ-0.37 2.0m FCM-HDC(19), Doric Lenses). The custom multi-fiber cannula used optical fibers of 200*µm* core, and borosilicate cladding resulting in an NA of 0.66 (HDFC 200/250-0.66 HD19 0002, manufactured by Doric Lenses). The spatial configuration of the fibers was designed such they would target the following locations (relative to bregma) in the mouse brain:

**Table 2.** Stereotactic coordinates of custom multi-fiber cannula targets. ^1^ mNAc: medial nucleus accumbens; vmNAc: ventral medial nucleus accumbens; vNAc: ventral nucleus accumbens; lNAc: lateral nucleus accumbens; DMSr, DMSi, DMSp, DMSc: rostral, intermediate, posterior and caudal parts of the dorsomedial striatum, respectively; DLSr, DLSi, DLSp, DLSc: rostral, intermediate, posterior and caudal parts of the dorsolateral striatum, respectively; TS: tail of striatum; TSp: posterior region of TS. *denotes the subset of fibers used in the photostimulation multi-fiber implant.

| Fiber number | 1* | 2 | 3* | 4* | 5* | 6 | 7 | 8 | 9* | 10 | 11 | 12 | 13 | 14 |
| Target region<sup>1</sup> | mNAc | vmNAc | vNAc | lNAc | DMSr | DMSi | DMSp | DLSr | DLSi | DLSp | DMSc | DLSc | TS | pTS |
|---|---|---|---|---|---|---|---|---|---|---|---|---|---|---|
| AP (mm) | 1.67 | 1.24 | 1.67 | 1.2 | 0.65 | 0.15 | -0.35 | 0.9 | 0.45 | -0.1 | -0.8 | -1.4 | -0.6 | -1.5 |
| ML (mm) | -0.6 | -0.85 | -1.2 | -1.74 | -1.65 | -1.95 | -2.35 | -2.35 | -2.7 | -3.0 | -2.7 | -3 | -3.35 | -3.65 |
| Fiber length v1 (mm) | 5.5 | 5.7 | 6.0 | 6.0 | 3.8 | 3.8 | 3.8 | 3.8 | 3.8 | 3.8 | 4.5 | 3.5 | 4.5 | 4.8 |
| Fiber length v2 (mm) | 5.2 | 5.4 | 5.7 | 5.8 | 3.8 | 3.8 | 3.8 | 3.8 | 3.8 | 3.8 | 4.5 | 3.5 | 4.5 | 4.6 |

Image acquisition was setup so that camera 1 collected the fluorescence emitted by GCaMP when excitated by LEDs 1 (410 *−* 420*nm*, isosbestic wavelength) and 2 (460 *−* 490*nm*, calcium-dependent modulation), while camera 2 collected fluorescence from tdTomato (excited by LED 3(555 *−* 570*nm*)). Each wavelength was acquired at 20*Hz* in a 640 *×* 640 pixel image. The objective of the BFMC6 was focused on the top FC connector of the optical fiber bundle. The cameras imaged the top of this fiber bundle where regions-of-interest (ROI) corresponding to each optical fiber were defined in the Doric Neuroscience Studio software (Doric Lenses). A 15*^th^* ROI was defined next to one of the fibers, to serve as a background control in the image. The average pixel value of each ROI was used as a measure of fluorescence emitted through each fiber (or control patch). The bundled fiber patchcord was connected to the custom fiber receptacle implanted in the mouse brain to deliver excitation light to the brain and collect the fluorescence emission signals from it simultaneously. The green and red fluorescence signals from the brain were spectrally separated from each other and from the excitation lights using dichroic mirrors integrated in the BFMC6. LED driving currents were chosen so that each excitation wavelength provided around 100*µW* at the tip of each optical fiber.

Synchronization with the behavioral data was achieved by receiving TTL pulses generated by the Bpod state machine in the photometry system, where each TTL pulse corresponded to one trial in the behavioral session.

#### Acquisition of electrophysiological data using Neuropixels probes

Four shank Neuropixels 2.0^87^ data was acquired through National Instruments PXI chassis (NI PXIe-1061 with NI PXIe-8381 and NI PCIe-8381) with an IMEC PXIe card (IMEC PXIE 1000). The software to acquire the data was SpikeGLX version 20190413-phase3B2 (https://billkarsh.github.io/SpikeGLX/). Pixels closest to the tip of the shanks were selected for recording in all four shanks. Synchronization with behavior was achieved through TTL pulses from the Bpod state machine (one pulse per trial) to the IMEC PXIe card and recorded through SpikeGLX. For these experiments, chronically implanted mice (with 1-5 fibers in the striatum) were prepared for acute recordings the evening before the first recording: a craniotomy was made above the VTA according to previously marked locations in the skull. The craniotomy was covered with a low toxicity silicon sealant (Kwik-cast, World Precision Instruments). The following day, the Neuropixels 2.0 probe for recording was rinsed in water and a dye was applied to its shanks (DiO, DiI, or DiD, Invitrogen, V22889, depending on the recording day).

Mice were placed on the head-fixed setup and their face was covered by a custom 3D-printed black case until the probe was inserted in the brain, to block their vision during preparations for probe insertion. The top of the head implants was cleaned and patchcord fiber(s) were connected to the implanted fibers’ receptacle(s). In the case of the custom multi-fiber implant (Extended Data Fig. 11a), we used a custom fan-shaped patchcord fiber bundle (HDP(5) 240/250/LWMJ/900-0.63 2.0m 5xFC-HD19 #2005, Doric Lenses) connected to 5 LEDs (CLED 465, Doric Lenses) at the opposite end from the implanted cannula. These LEDs were controlled by a single LED driver (LEDD 8, Doric Lenses) which in turn was triggered by the Bpod state machine and its Analog Output Module (1018, Sanworks). For the 3-fiber and 1-fiber configurations (Extended Data Fig. 11f,k) we used a single patchcord fiber (MFP 400/440/110 −0.37 2m FC-ZH1.25(F) LAF, Doric Lenses) for each implanted cannula fiber, each connected directly to a laser (LRS-0473-GFM-00100-03 with PSU-III-LED power supply, Laserglow Technologies).

The craniotomy cover was removed and the brain surface was kept covered with saline. The Neuropixels probe was connected to the acquisition system and aligned to the insertion area in the craniotomy using a micromanipulator (PTC1-Z8, Thorlabs). The probe was inserted in the brain until reaching the recording region (VTA or SNc) with a velocity profile of 0.1*mm/s*, 0.01*mm/s*, 0.005*mm/s*, 0.002*mm/s*, 0.0001*mm/s* with transitions at 0.5*mm*, 2*mm*, 3*mm*, 3.5*mm* from the brain surface. The probe was inserted 100*µm* deeper than the target, retracted to the target coordinate, and then kept stable for 10 to 30 minuted before recording started. Recording of neural data was performed using SpikeGLX (https://billkarsh.github.io/SpikeGLX/) while mice performed the behavioral task, during the optotagging protocol, and during the air puff delivery protocol (which was used in a subset of the mice). After the last behavioral protocol was finished, the Neuropixels probe was removed by moving it towards the brain surface at a velocity profile of 0.002*mm/s*, 0.005*mm/s*, 0.01*mm/s*, with transitions at 1*mm*, and 1.5*mm* above the recording site. After removal, the probe was dipped in distilled water for 20 min before being left in 1% terg-a-zyme (Z742918, Sigma-Aldrich) overnight. The patchcord fiber(s) were detached from the implanted fiber receptacle(s), the craniotomy was washed, the skull dried, and the silicon sealant was used to cover the craniotomy. Acute recordings were performed in the same mouse for 4 or 5 consecutive days, while varying slightly the location of the probe penetration in the brain.

#### Optotagging protocol

Similar to behavioral protocols, the optotagging protocol was written in Matlab and controlled through the Bpod state machine, which communicated with an analog output module (1018 or 1013, Sanworks) to trigger light pulses from LEDs or laser(s).

In the case of the 5-fiber multi-fiber implant (Extended Data Fig. 11a), 5 LEDs (CLED 465, Doric Lenses) were controlled by a single LED driver (LEDD 8, Doric Lenses) to deliver light through each one of the fibers in the fiber bundle (HDP(5) 240/250/LWMJ/900-0.63 2.0m 5xFC-HDC(5) 0005, Doric Lenses) connected to the implanted cannula. The protocol consisted in the delivery of a total of 300 trials, 50 trials for each one of the 6 trial types: one trial type corresponded to deliver of light from one of the optical fibers, and the last trial type consisted in simultaneous illumination of all optical fibers. In each trial, 10 pulses of light with a duration of 30*ms* at 10*Hz* was delivered, resulting in a total of 500 delivered pulses per optical fiber. The inter-trial-interval (ITI) between trials was taken from an exponential distribution between 2*s* and 4*s*.

In the case of the 3-fiber or 1-fiber antidromic optotagging configuration (Extended Data Fig. 11f,k), the analog output triggered diode-pumped lasers directly connected to an optical fiber patchcord each (MFP 400/440/110-0.37 2m FC-ZH1.25(F) LAF, Doric Lenses). The photostimulation/optotagging protocol consisted in the delivery of 200 trials per laser, 10 pulses per trial, each with a duration of 20*ms* delivered at 10*Hz*, using an ITI exponential distribution between 0.5*s* and 3*s*.

#### Histology and Immunohistochemistry

After the last experiment, mice (except multi-fiber photometry mice, see below) were deeply anesthetized with an overdose of ketamine/medetomidine, exsanguinated with 0.9% phosphate buffered saline (PBS), and transcardially perfused with cold 4% paraformaldehyde (PFA) in PBS. The brain was removed from the skull and kept in PFA for 24 hours at 4*^◦^C* and then rinsed and stored in PBS at 4*^◦^C* until slicing. Brain slicing was performed in a vibratome (VT1000S, Leica), to obtain sections of 70 or 100*µm*. Slices were either directly mounted on glass slides using a mounting medium with 4’,6’-diamidino-2-phenylindone (DAPI, VECTASHIELD, Vector Laboratories), or prepared for immunostaining with tyrosine hydroxylase antibody (AB152, Sigma Aldrich) to visualize dopamine axons or cell bodies before mounting. Mounted brain slices were imaged with an automated fluorescence microscope (Axio Scan.Z1, Zeiss) or in the confocal microscope (LSM 700, Zeiss). The locations of the tracks of each Neuropixel shank were annotated in relationship to the midline of the brain slice and the tips of the shanks in relation to the ventral surface of the brain in the Zeiss Zen Blue software.

#### Tissue fixation and staining for micro-CT

Mice implanted with the custom 14-fiber implant for multi-fiber photometry recordings were deeply anesthetized with isofluorane and exsanguinated with artificial cerebro-spinal fluid (ACSF) for 2-3 minutes at 10*ml/min* and then transcardially perfused with a solution of 2% PFA and 2.5% glutaraldehyde (16220, Electron Microscopy Sciences) in PBS. The brain was fully separated from the skull in some cases, and in other cases the dorsal skull was kept attached to the brain with the multi-fiber implant still implanted in the brain. The brains (with or without implants attached) were kept in the same fixative solution (2% PFA and 2.5% glutaraldehyde) at 4*^◦^C* for 48 hours, and then rinsed with PBS at 4*^◦^C* for 24 hours. Brains were stained with 2% osmium tetroxide (19190, Electron Microscopy Sciences) in sodium cacodylate buffer (11650, Electron Microscopy Sciences) in an orbital shaker at room temperature for 2 *−* 4 days. The solution was then switched to PBS for at least 24 hours at 4°*C* before scanning.

#### Acquisition of micro-CT images

X-ray micro-CT was performed using a Zeiss Xradia 520 Versa X-ray microscope. The samples were wrapped in gauze to ensure stability during scanning and subsequently immersed in PBS in a polystyrene jar (PLC03356, Qorpak) with the cap tightly sealed. The jar was then secured to the rotary stage in the X-ray recording chamber using double-sided adhesive tape. Scans were acquired using the Scout-and-Scan Control System (Zeiss). Prior to each scan, the sample position was adjusted to the center of the field of view and the imaging protocol was set up. The sample was imaged using a 0.4*×* objective lens at a tube voltage of 60*kV* and an output power of 5*W*. The exposure time was set at 2.5*s*. For each tomographic scan, the sample was rotated 360*^◦^*, and either 2401 or 2201 projection images were acquired over one full rotation.

### 5.2 Data curation and analysis

Data curation, preprocessing, analysis (including statistical tests), and computational modeling were performed using custom code in MATLAB (versions 2017a, 2019b, 2025b) and Python (versions 2.7.18, 3.11, 3.14.4) using packages NumPy, SciPy, pandas, scikit-learn, statsmodels, pyGAM, pynapple, matplotlib, and seaborn.

#### Multi-fiber photometry data

In the freely behaving setting (Extended Data Fig. 1) the average pixel value of each fluorescence output of each ROI (14 optical fibers + 1 control ROI) was preprocessed by calculating Δ*F*/*F*_0_ as 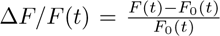 using as *F*_0_ the 6*^th^* running percentile in a window of 40*s*. This Δ*F/F*_0_ was then normalized to its maximum, and the cross-correlogram was obtained for all pairs of fibers. The auto-correlogram of each fiber was fit with an exponential function of the form 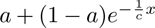 to obtain the autocorrelation decay time constant, *c* (Extended Data Fig. 1d,e). The pairwise Pearson’s correlation coefficients were calculated for each fiber pair of each mouse, and then averaged across mice for each striatal target. The average cross-correlation matrix was used to create a hierarchical binary cluster tree using the linkage function in Matlab.

To analyse fiber-photometry data in the head-fixed behavioral tasks, average pixel value of each fluorescence output of each ROI was first aligned to the time of odor delivery. Δ*F/F*_0_ was calculated considering *F*_0_ the average signal during 1*s* prior to odor delivery. Possible movement artifacts in the functional channel were corrected by subtracting from the GCaMP Δ*F/F*_0_ trace a prediction of movement signal calculated using linear regression^88^ from either the isosbestic or the tdTomato Δ*F/F*_0_ signal, whichever had the most significant correlation coefficient to the GCaMP non-corrected Δ*F/F*_0_. This movement corrected signal was then z-scored across the entire session (separately for each fiber). Quantification of cue and outcome responses was done by averaging the movement-corrected z-scored Δ*F/F*_0_ GCaMP signal in the time intervals 100 *−* 800*ms* and 300 *−* 800*ms* following cue initiation and reward initiation, respectively.

For the analysis of temporal shift^33^, we searched in the period between the cue and the reward delivery for activity peaks 3*×* larger than the standard deviation of the baseline period and chose the largest if there were more than one. Delay to the peak (from cue initiation) was plotted as a function of trial number during learning and a regression slope was obtained to evaluate the presence of temporal shift of this peak over trials. The same type of linear regression was obtained for evaluating a change in cue response amplitude, and in the amplitude to a fixed reward as a function of trial number (Extended Data Fig. 5, 5). These quantifications of calcium fluorescence signal were used to build the RPE index as described in the main text.

#### Preprocessing of micro-CT images

Micro-CT volumes were converted from the Zeiss TXM file format to NRRD format using custom-written Python code (https://github.com/YuelongWu/xradia_scripts). A 3x3x3 median filter was applied to all volumes to suppress noise. The grayscale values of the volumes were scaled based on the number of projections in the tomography data to maintain consistency. To register the volumes into a common reference frame, they were first manually aligned to an approximate initial orientation, followed by registration refinement using Elastix software^89,90^ (https://elastix.dev/) with a Euler transformation. The registered brain scans were then registered to the Allen Mouse Brain Common Coordinate System version 3 (CCFv3) using affine transformation in Elastix. Atlas and annotation volumes were obtained from the Scalable Brain Atlas *ABA_v_*3^91^ implementation of the Allen Mouse Brain Atlas^92^ and manual labeling of optical fiber tip coordinates was done in FIJI^93^.

### Preprocessing of Neuropixels data

Neuropixels data was spike-sorted using Kilosort3 integrated in the ecephys pipeline (https://github.com/jenniferColonell/ecephys_spike_sorting by Jennifer Colonell, Allen Institute). This pipeline uses CatGT (https://github.com/billkarsh/CatGT) for high-pass filtering of the raw data and removal of artifacts before spike sorting, extracts synchronization pulses, and establishes quality metrics after spike-sorting. Manual curation of the Kilosort3 output was done with Phy (https://github.com/cortex-lab/phy), followed by re-calculation of unit quality metrics using the ecephys pipeline again. Finally, spike times were synchronized with behavioral data from Bpod, followed by the optotagging tests (see below).

#### Peristimulus time histogram

Taster plots and peristimulus time histograms (PSTHs) of neural activity in spiking activity were computed using a MATLAB toolbox written by HyungGoo Kim (https://github.com/HRKimLab/libkm). PSTHs were constructed using 1-ms bins and the signal was not smoothed during preprocessing (e.g. when histograms were used for photoidentification of dopamine neurons). PSTHs used for distributional RL data analysis were smoothed by convolving with the function *f* (*t*) = (1 *− e^−^t*)*.e^−t/τ^*, where *τ* is a time constant set to 20*ms* as in^2,60^. For plotting purposes, firing-rate traces were further smoothed by convolution with a unit-area Gaussian kernel (1*s* window, *σ* = 33*ms* for population traces and *σ* = 45*ms* for single-neuron traces). For summary plots

#### Antidromic identification of dopamine neurons

We aimed to detect antidromic propagation^94^ of light-triggered action potentials in the cell bodies of dopamine neurons expressing ChR2 following light stimulation of their axons in the striatum^34^. The criteria for identification of antidromically tagged dopamine neurons were established by pooling together all Neuropixel recording sessions, independently of behavioral task, or of the configuration/number of optical fibers implanted in the striatum (Extended Data Fig. 11). Identified dopamine neurons had to satisfy all of the following criteria, for any individual optogenetic stimulation target in the striatum:

1. satisfy the quality metrics requirements recommended by the Allen Institute at the output of the ecephys pipeline: *presence ratio >* 0.9, *isi viol <* 0.5, *amplitude cutoff <* 0.1;
2. the firing rate in any 1*ms* bin within a 20*ms* time window following the light onset had to be at least 5 standard deviations above the mean of the shuffled firing rate;
3. the Pearson’s correlation between the mean spontaneous and the mean lighttriggered spike waveforms had to be above 0.9;
4. the stimulus-associated spike latency test (SALT)^95–97^ had to be passed with a pvalue lower than 0.05, using 1*ms* bins and a test window of 20*ms* after the light onset, in all of these three conditions: a) at least for one of the 10 pulses delivered within a trial (concatenation across trials but not across pulses within single trials) using as null distribution the latencies taken from 20*ms* time windows in a 1.4*s* interval within the ITI; b) when all light pulses were concatenated (i.e. without regard to their presentation order within the trials) using as null distribution the last 30*ms* before the light pulses; and c) same as b) while removing all laser pulses that corresponded to the second laser pulse in a trial (which occurs at 100 ms after light pulse train onset - this was done because we often saw variable latency spike activations around this delay, which we presume is light-evoked activity triggered with the start of the light pulse train, Extended Data Fig. 11g,l);
5. it could not violate the action-potential collision test: which consisted of selecting the individual light pulses that coincided with spontaneous spikes. These were spikes that occurred between *−*7 and *−*2*ms* before the expected latency for the light-triggered response, defined as the first two 1*ms* time bins in which the firing rate first passed the shuffle test (point 2)). The firing rate of these stimulation trials at the expected latency for the laser-triggered response had to be zero to pass this test (in either one of these two 1*ms* bins). This collision test acted as an exclusion test, to remove units that violated it. We note, however, that some neurons had a low baseline firing rate (*n* = 18 out of *n* = 323 optotagged neurons) resulting in no trials available to make this test an inclusion test, precluding a more thorough statistical collision test.

#### Analysis of single unit recordings

Single unit PSTHs were baseline-corrected for distributional RL analysis, but not for receiver operating characteristic curve (ROC) or for MLE analysis. The baseline correction consisted of subtracting, from each trial, the mean over the activity of that trial from 1 to 0*s* relative to the cue onset. Plots with average single neuron activity along each trial type were obtained by computing a time-resolved area under the receiver operating characteristic curve (auROC). Spike density functions were first averaged into 50*ms* bins. For each neuron, the firing-rate distribution in each time bin was compared with the distribution of firing rates during the 900*ms* baseline period preceding cue onset by sweeping a firing-rate criterion across the full range of observed firing rate for the neuron. The area under the ROC curve was computed by numerical integration, yielding a time-resolved auROC value for each bin. An auROC value of 0.5 indicates no difference between the baseline and test distributions, whereas values greater or smaller than 0.5 indicate increases or decreases in firing rate relative to baseline, respectively. For population averages, trial type firing rates were averaged across all neurons projecting to a particular striatal target. Reward-evoked responses were defined as the average activity from 200 to 600*ms* after reward onset as in previous studies^2^.

#### Empirical estimation of distributional RL parameters (standard fitting procedure)

To estimate the parameters underlying distributional RL from single dopamine neurons, we used the method developed by Dabney et al., 2020^2^. We quantified two quantities for each neuron: the zero-crossing point, corresponding to the reward magnitude that elicited no prediction-error response, and the optimism level (*τ*), which quantifies the relative weighting of positive and negative prediction errors. Estimating both quantities from the same trials can introduce statistical dependencies between them. To obtain independent estimates, we randomly partitioned the trials of each reward size into two non-overlapping halves. One half was used exclusively to estimate the zero-crossing point, whereas the other half was used to estimate the slopes of positive and negative reward responses. A common utility axis was first constructed from the average reward response of each projection-defined population as a function of reward size, here referred to as the utility function following work from Stauffer and colleagues^2,68,98^. The population-average reward response defining the utility axis was smoothed using a monotonically increasing spline (SLM toolbox, MATLAB) to obtain a strictly monotonic utility function. This procedure minimizes the effects of measurement noise while preserving the expected monotonic relationship between reward magnitude and utility, allowing reliable interpolation between utility space and reward magnitude. For each neuron, candidate zero-crossing locations were defined halfway between consecutive reward magnitudes. For each candidate location, responses to reward sizes above the candidate were classified as positive prediction errors and responses to reward sizes below the candidate as negative prediction errors. A criterion score was then calculated as the total number of responses consistent with this classification (i.e., positive responses above the candidate and negative responses below it). The zero-crossing point was defined as the candidate location maximizing this criterion. Small random perturbations were added to the criterion values to avoid ties between adjacent candidates. Zero-crossing estimation was performed in utility space (i.e. from the firing rate), the discrete estimate was then converted into reward magnitude using the continuous utility function constructed for the population (as described above). Zero-crossing in reward magnitude space was obtained by weighted interpolation between the neighboring points of the utility function. The utility axis was then shifted by the estimated zero-crossing to center each neuron’s response function before estimating the positive (*α*^+^) and negative (*α^−^*) learning rates by piecewise linear regression on either side of the zero-crossing. The optimism level was calculated as 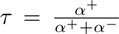. Neurons with undefined or with negative estimates of either scaling factor were excluded from subsequent analyses, as they violate the monotonic reward-response relationship assumed by the distributional RL framework. This random partitioning and estimation procedure was repeated 50 independent times: for each partition, correlations between zero-crossing estimates and optimism level obtained from the two halves of the data were computed and displayed in a histogram (Extended Data Fig. 13). Final estimates of zero-crossing and optimism for each neuron were obtained by averaging across iterations (Fi.4i).

The same analysis was repeated using as utility function the global average reward response as a function of reward size (i.e. all dopamine neurons were pooled together for this global utility function, independently of projection target). Results obtained were qualitatively similar (Extended Data Fig. 14).

#### Distribution decoding from dopamine populations

Reward distributions were reconstructed using the expectile decoding algorithm of Dabney et al. (2020), implemented using truncated Newton constrained optimization (TNC)^2^. We performed decoding from simulated value predictors and from each projection-defined dopamine population. Decoding was run both following empirically estimated zero-crossing points and optimism levels, as well as following MLE estimation of zero-crossing points and optimism levels. Because the VS+DS-projecting population contained the smallest number of neurons (11), in Figure 4 all other projectiondefined populations were repeatedly subsampled without replacement to the same number of neurons. One thousand random subsets were generated for each population, whereas ten decoding runs were performed for the VS+DS-projecting population using the complete dataset. As a control, optimism levels and zero-crossing points of VS+DS-projecting neurons were randomly permuted prior to decoding, disrupting their correspondence while preserving the marginal distributions of both variables.

Decoding performance was quantified using three complementary measures. First, the decoding loss returned by the optimization algorithm was used to evaluate the quality of the reconstructed distribution. Second, similarity between decoded and empirical reward distributions was quantified using the one-dimensional Wasserstein distance (Earth Mover’s Distance). For each projection-defined population, Wasser-stein distances were computed between the empirical reward distribution and each decoded distribution. When multiple decoding runs were available, the lowest-loss decoding was selected for shaded plotting in Figure 4f. Mean Wasserstein distances and 95% confidence intervals were estimated by bootstrap resampling. Finally, we assessed whether experimentally estimated zero-crossing points matched the theoretical expectile values predicted from their optimism levels. For each neuron, the empirical reward distribution was used to determine the theoretical expectile corresponding to its optimism level, and the agreement between predicted expectile values and experimentally measured zero-crossing points was quantified using the coefficient of determination (*R*^2^).

### 5.3 Computational modeling

#### Simulation of distributional reinforcement learning

We simulated distributional RL using a population of value predictors trained with a Rescorla–Wagner learning rule implementing asymmetric learning rates. Simulations were performed using 11 value predictors, matching the number of dopamine neurons available for decoding from the experimentally identified VS+DS population. For each predictor *V_i_*, positive and negative reward prediction errors were weighted by asymmetric learning rates 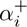and 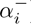), resulting in an optimism level 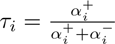 Learning rates were randomly sampled while maintaining a constant total learning rate 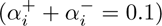 producing a population spanning the full optimism spectrum and showing convergence in 300 *−* 500 trials. Simulations were run on multiple reward distributions, namely the one used in the behavioral task that mice were trained on, and value predictors converged to the corresponding expectiles of the reward distributions. Following learning, the reward distribution was reconstructed from the simulated expectile representation using the expectile decoding algorithm described by Dabney et al. (2020)^2^. Ten independent simulations were performed for each reward distribution.

#### Maximum likelihood estimation for zero-crossing points and optimism

For each identified dopamine neuron, we modeled its trial-by-trial reward response as a linear function of reward magnitude, following the distributional RL algorithm (Fig. 1b). Specifically, we fit

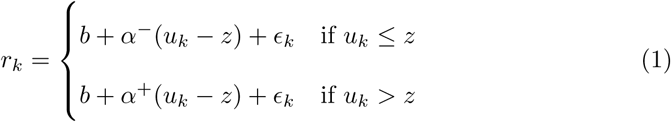

where *z* is the zero crossing point, *α^−^* and *α*^+^ are the slopes for the negative and positive prediction errors, respectively, and *ɛ_k_ ∼ N* (0*, σ*^2^) with *σ* = 2Hz. The response on trial *k*, *r_k_*, was the mean firing rate in the 200*−*600*ms* window after reward delivery, and the neuron’s baseline *b* was its mean firing rate in the 1*s* preceding cue onset, averaged across all trials. Reward magnitudes were first mapped onto a utility axis, *u*(*R*), given by the population-average, baseline-subtracted response as a function of reward size, and smoothed with a monotonically increasing spline. Finally, we used a Gaussian observation model, where

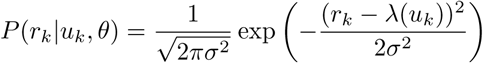

The baseline and responses were set to the empirical values such that only *θ* = (*z, α^−^, α*^+^) were fit. The three parameters were estimated jointly for each neuron by minimizing the negative log-likelihood,

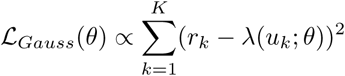

using the L-BFGS-B algorithm, with bounds *z ∈* [*−*10, 25] on the utility axis and *α*^+^*, α^−^ ≥* 0. Trials with zero spikes were assigned a small positive rate randomly drawn from *|N* (0, 0.001) to avoid degenerate likelihoods. Optimization was initialized from the empirical estimates of zero-crossing and slopes obtained with the procedure from^2^; neurons for which those estimates were undefined were initialized from the population mean.

A neuron was considered “not fitted” or “lost” (Fig. 3j, 4c) if the optimizer failed to converge, if either slope was undefined, or if either the fitted slope was zero or negative (i.e., the response was not monotonically increasing with reward on both sides of the zero-crossing, as required for canonical RPE coding). For neurons that were successfully fit, the optimism level was calculated as 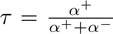 and the zero-crossing point was converted from the utility axis back to reward volume (*µ*L) by linear interpolation of the population response function, extended by one step at each end to allow zero-crossings outside the range of delivered rewards. Estimates in these regions are poorly constrained and the decoder we use confines samples to the reward range, so neurons with zero-crossings in this region will inflate decoding loss for that population.

MLE estimation was validated against the original MATLAB implementation of Dabney et al. 2020^2^ on their published data set^60^.

#### Confidence intervals

Uncertainty on each neuron’s MLE parameters was estimated by non-parametric bootstrap and shown in Extended Data Figs. 15, 18. Trials were resampled with replacement within each reward magnitude, the model was refitted to the resampled data (initialized at the original estimates), and this was repeated *B* = 1000 times. 95% confidence intervals were the 2.5th and 97.5th percentiles of the bootstrap distribution of *z* (after conversion to *µ*L) and of *τ*.

The correlation between zero-crossing and optimism across neurons (Figs. 15, 18D) was assessed in two ways. First, to quantify sensitivity to the particular neurons sampled, neurons were resampled with replacement *B* times and the Pearson correlation recomputed on each resample. Second, to quantify sensitivity to trial-to-trial noise in the fits, the *b*-th bootstrap fit of every neuron was combined into one population and the correlation computed across neurons, for each fit. In both cases 95% confidence intervals were taken from the 2.5th and 97.5th percentiles.

#### Decoding the reward distribution from MLE estimates

The reward distribution encoded by each population was decoded from the fitted (*z, τ*) pairs with the expectile decoder of Dabney et al.^2^: a set of *N* = *n −* 1 samples (with *n* the number of fitted neurons) was optimized so that its *τ* -expectiles matched the zero-crossing points *z* (in *µ*L), by minimizing the squared expectile loss with bounded optimization (truncated Newton method; samples restricted to [0, 20.1] *µ*L). The optimization was repeated over 10 epochs from 50,000 random initializations each, and the resulting samples pooled.

Decoding quality was quantified by the 1-Wasserstein distance between the decoded samples and 1,000 samples drawn from the true reward distribution, and by the Kullback–Leibler divergence, KL(true ‖decoded), between Gaussian kernel density estimates of the two evaluated on a 500-point grid. Results are shown in Fig. 15e and Fig. 18e. Because the decoder is stochastic, both metrics are reported as mean *±* SD over 10 independent decoding runs.

### 5.4 Data availability

Preprocessed data are documented and will be posted to an online repository upon publication.

### 5.5 Code availability

The code used for analysis, computational modeling, and generation of figures will be posted to an online repository upon publication.

## 6 Acknowledgements

We thank HyungGoo Kim for help setting up the antidromic optotagging recording technique, Ryunosuke Amo for technical advice, Paul Masset for comments on the manuscript, and all members of the Uchida laboratory for valuable discussions and comments on the project. We thank Edward Soucy, Brett Graham, and Yuwei Li of the Neurotechnology Core for providing critical assistance with instrumentation, and the Harvard Center for Biological Imaging (RRID: SCR 018673) for infrastructure and support for *ex vivo* imaging. This work was supported by grants from the US National Institutes of Health (R01NS116753 to N.U. and J.D., 5R01DA05975 to N.U. and M.W.-U., F31NS124095 to A.S.L., and 1K99DA060290 to M.C.), the Simons Collaboration on Global Brain (to N.U.), the Human Frontier Science Program (LT000801/2018 to S.M.), the Harvard Brain Science Initiative, and the Brain and Behavior Research Foundation (NARSAD Young Investigator no. 30035 to S.M.).

## 7 Author contributions

S.M. and N.U. designed the experiments. S.M. and S.X. performed the experiments and preprocessed the data. M.C. established Neuropixels-based acute recording with antidromic optotagging with help from HyungGoo Kim. S.M. developed multi-fiber implants for fiber photometry and antidromic optotagging. S.M., S.X., X.L. developed micro-CT protocols under the supervision of J.W.L.. R.S. and Y.W. acquired and preprocessed micro-CT scans, respectively. A.L. implemented simulations and computational tools. C.R.B. developed the Maximum Likelihood Estimation procedure under the supervision of J.D. S.M. analyzed the data, implemented computational models and simulations, wrote the first draft of the manuscript, and created the figures. M.W.-U, J.D., and N.U. supervised the project. All authors edited the manuscript.

## 9 Extended data figures and legends

**Extended Data Figure 1.**
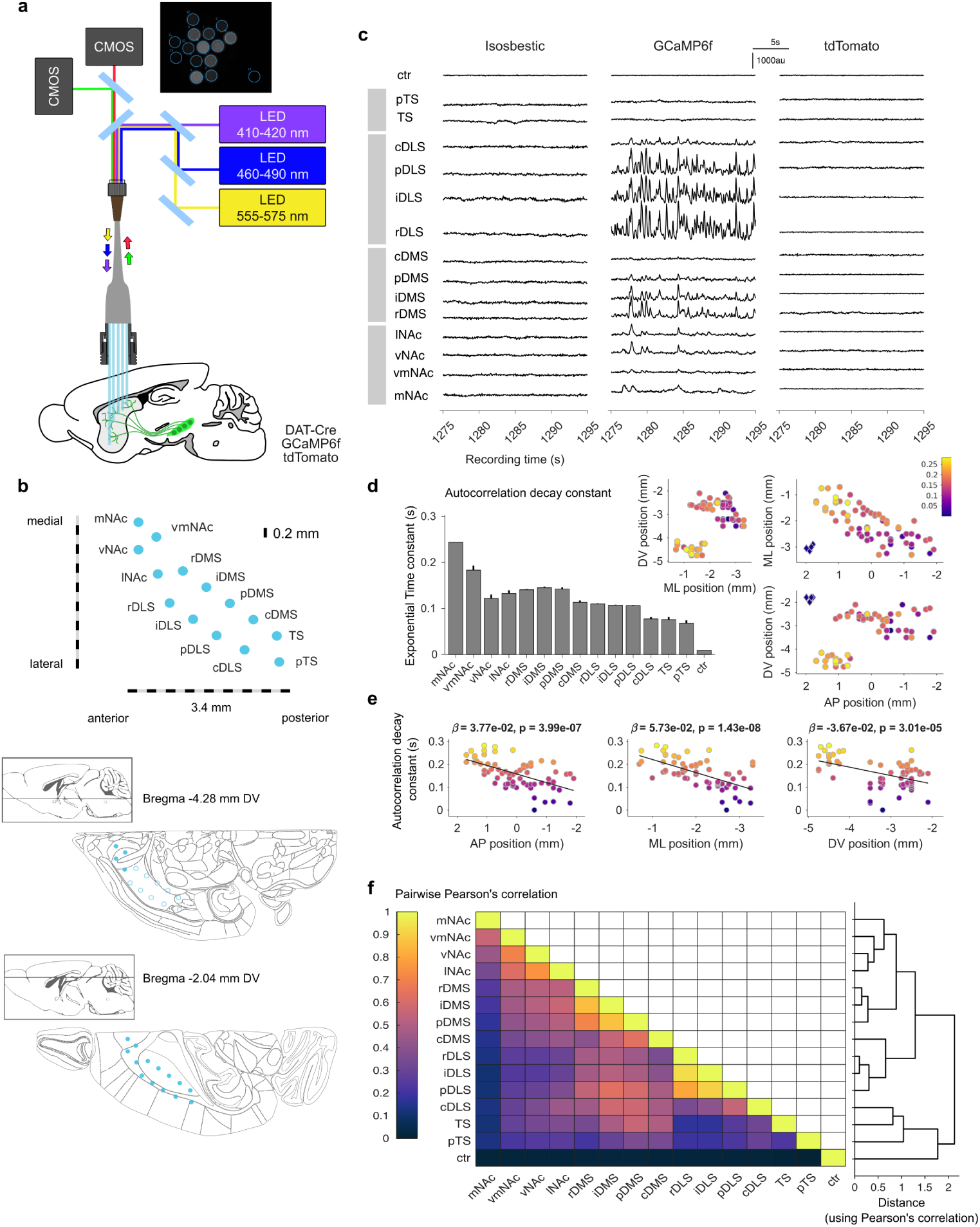
Multi-fiber photometry recording in behaving mice. **a**, Schematic of recording setup using a multi-fiber implant, and example field of view of one of the CMOS cameras. **b**, Top: optical fiber configuration of multi-fiber implant on viewed from the top horizontal plane with target region information: mNAc medial nucleus accumbens, vmNAc ventro-medial NAc, vNAc ventral NAc, lNAc lateral NAc, DMS dorsomedial striatum, DLS dorsolateral striatum, r rostral, i intermediate, p posterior, c caudal, TS tail of the striatum. Bottom: representation of the optical fibers in two example horizontal slices of the mouse brain: the ventral slice is the target of the NAc fibers (filled circles) but it is not reached by the dorsal fibers (open circles); the dorsal slice is traversed by all fibers. **c**, Example raw fluorescence traces acquired simultaneously for all 15 ROIs (14 in the brain and one control ROI not connected to any fiber) during alternating isosbestic, GCaMP6f, and tdTomato excitation cycles (effective acquisition rate for each channel is 20Hz). These example traces were acquired during free behavior. **d**, Exponential time constant after fitting to the autocorrelation plot of each ROI; left: averages of eleven mice across fiber targets during free behavior; right: autocorrelation time constant of each fiber in its final target location pooled across 5 mice in a freely behaving setting (diamonds represented in coordinates (2*mmAP, −*3*mmML, −*2*mmDV* are the control ROIs, for reference). **e**, Autocorrelation decay constant as a function of anatomical position in the anterior-posterior (AP), medio-lateral (ML), and DV (dorso-ventral) axis in the freely behaving setting. A significant gradient is observed across the three anatomical axes. **f**, Pair-wise Pearson’s correlation coefficients calculated on the GCaMP6f signal in a freely behaving setting (left), and corresponding dendogram (right), recovering the macro-level organization of the striatum characterized by topographic cortico-striatal projections into NAc, DMS, DLS, and TS, but also demonstrating complexity within these regions (e.g. notice how mNAc is different from other NAc regions).

**Extended Data Figure 2.**
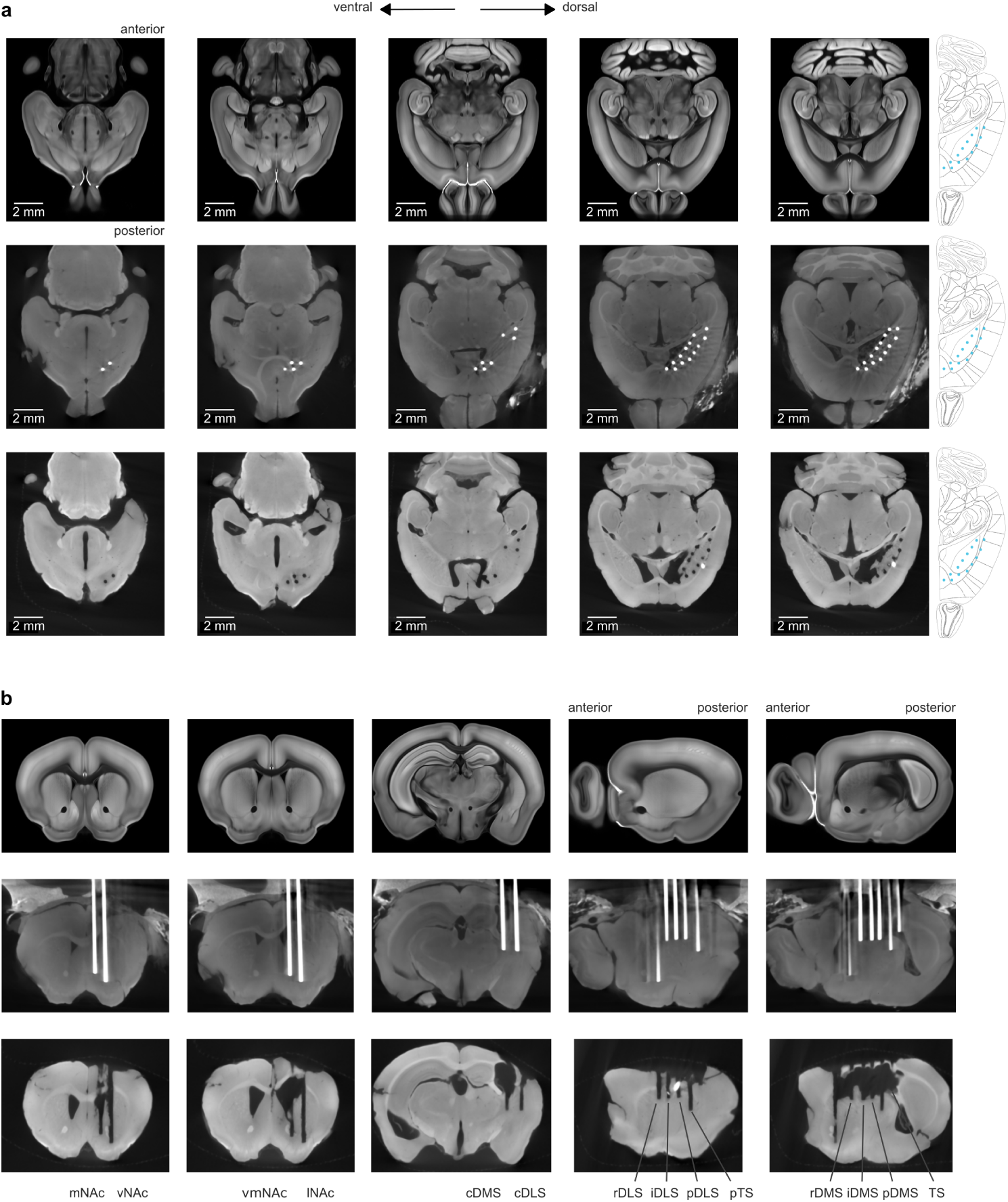
Micro-CT for fiber location assessment. **a**, Horizonal slices of the Allen Mouse Brain Atlas Coordinate Framework version 3 (top row) and corresponding horizontal slices from the micro-CT scan of the brains of two example mice: one in which the fiber implant was not removed prior to scanning (middle row) and one in which it was removed (bottom row). **b**, Same brains as above sectioned to reveal different views of fiber placements: coronal slices to show the NAc fibers (first two columns), and diagonal slices to show the caudal DMS and DLS fibers (middle) as well as the DLS and DMS fibers (two rightmost columns. Both preparations (keeping or removing the fiber implant) show fiber locations, but each one has its drawbacks: the presence of fibers introduces artifacts in the micro-CT images decreasing contrast quality, while removing the implant can cause further damage to the dorsal brain.

**Extended Data Figure 3.**
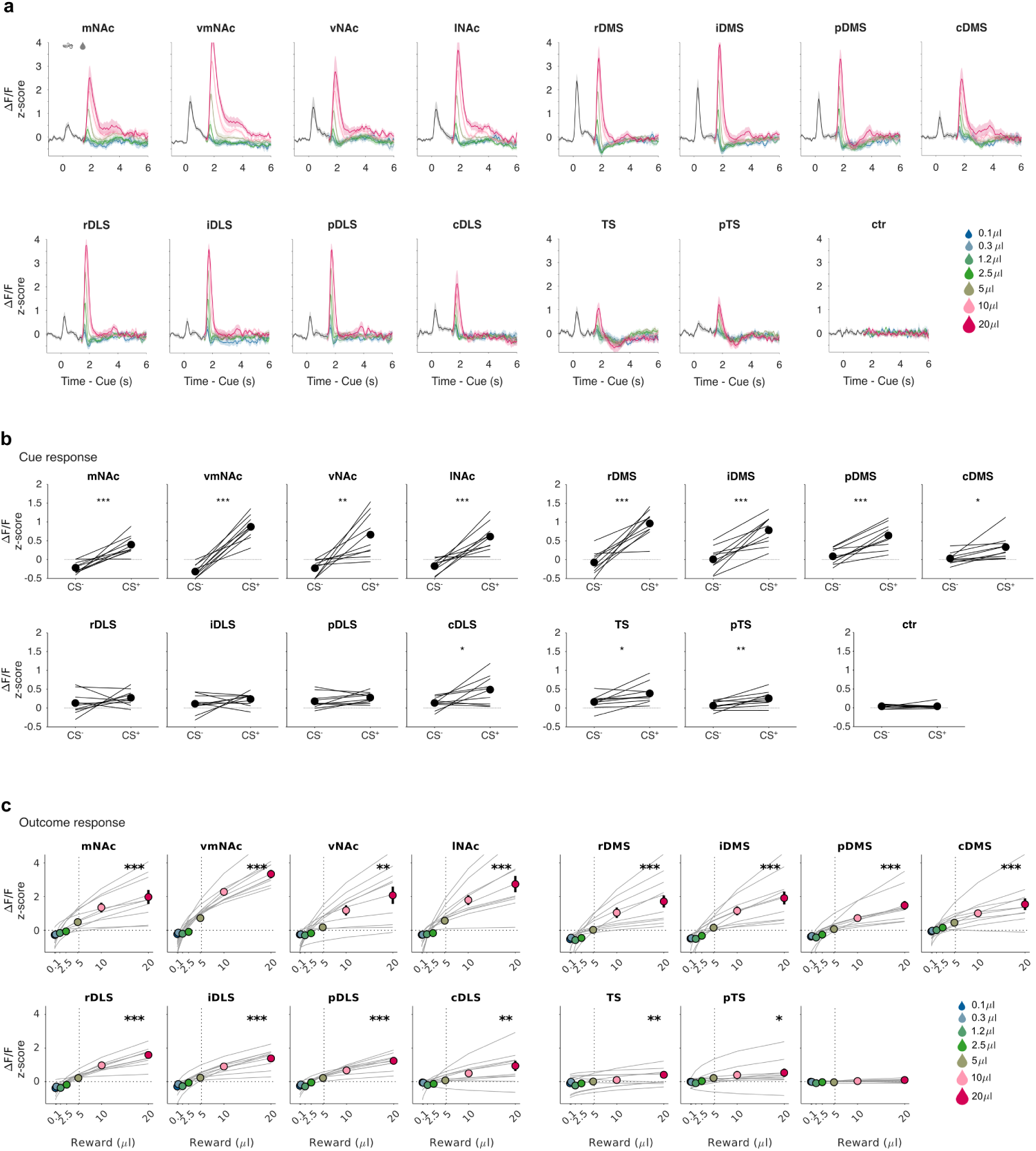
Cue and outcome responses across striatal regions. **a**, Average fluorescence traces across mice (*n* = 11) for each projection target and color coded according to reward size delivered 1.5*s* after odor initiation (time before reward is average of all *CS*^+^ trials, shaded areas represent standard error of the mean). **b**, Average cue response to *CS*^+^ and *CS^−^* odor cues (lines represent individual mice). Asterisks denote significance of paired t-test comparing *CS*^+^ and *CS^−^* responses. **c**, Average outcome response to each reward size delivered in *CS*^+^ trials. Colored circles indicate mean ± s.e.m. of experimentally measured dopamine axonal responses across mice; grey lines represent individual mouse response functions fitted with Hill model (or a linear model in case the Hill model fails to fit the data). Positive modulation by reward magnitude was assessed by fitting a linear regression to the responses of each mouse and testing whether the mean slope across mice exceeded zero (one-sided one-sample t-test). P-values were corrected across recording sites using the Benjamini–Hochberg false discovery rate procedure. (*** *P <* 0.001, \*\**P <* 0.01,\**P <* 0.05).

**Extended Data Figure 4.**
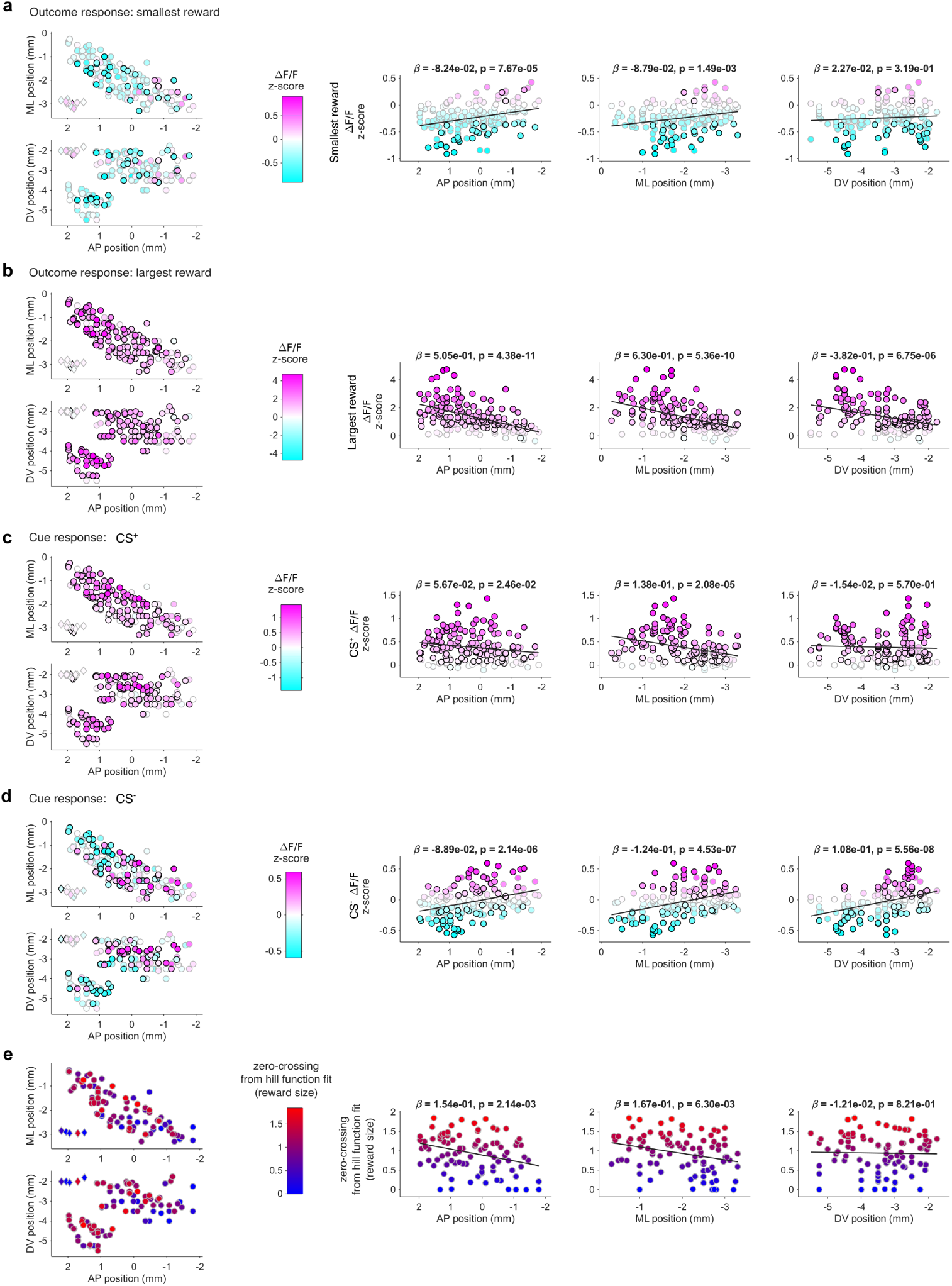
Cue and outcome responses across striatal regions: individual fibers and anatomical gradients. **a**, Average fluorescence response (z-scored Δ*F/F*_0_) to the smallest reward measured in each fiber (14 fibers in each of *n* = 11 mice, (same data as in Fig. 2d). Left: data shown in the AP-ML projection (top, horizonal view) and AP-DV projection (bottom, sagittal view). Diamonds plotted around the (2*AP, −*3*ML, −*2*DV*) coordinates correspond the control ROIs in the cameras’ fields of view, not to real fibers in the brain. Fibers with black edge show significant modulation from baseline (*P <* 0.05 Wilcoxon sign rank test), fibers with grey edge signal non-significant modulation from baseline. Right: Same data shown in each anatomical axis with slope (*β*) and significance value of a linear fit of the data to each anatomical axis. **b**, same as **a** for the largest reward delivery. **c**, same as **a** for *CS*^+^ response. **d**, same as **a** for *CS^−^* response. **e**, Same as **a** for the inferred zero-crossing point after fitting a hill function to the reward response function of each fiber. Only the zero-crossing point of successful fits are shown. A significant gradient is observed in the medio-lateral axis as previously reported^49^, but the range of zero-crossing values is limited and close to zero, as fiber photometry recording of calcium activity have a smaller range of activity for decreases in neural activity than to increases^99^.

**Extended Data Figure 5.**
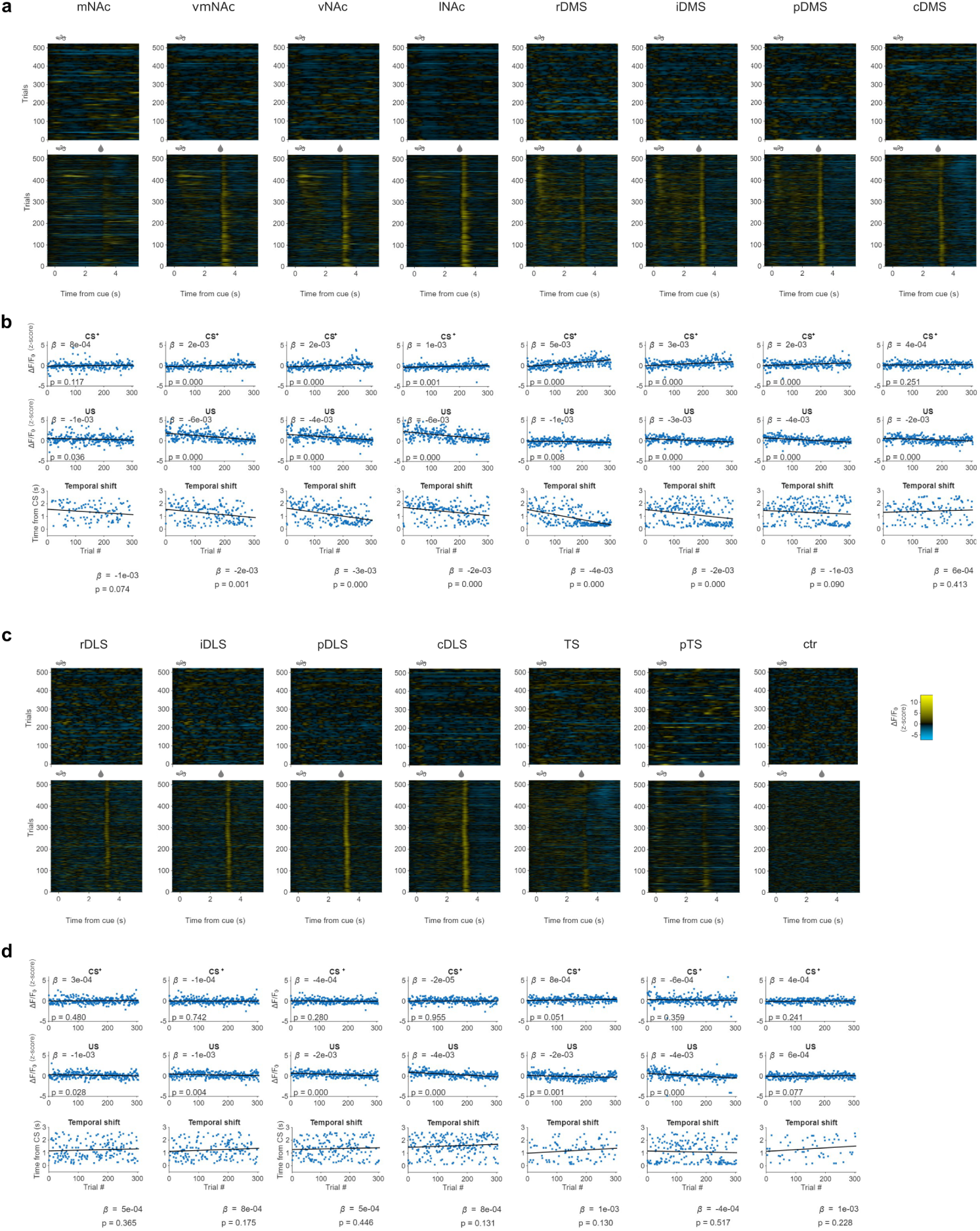
Modulation of cue and outcome responses during learning across striatal regions: example mouse. **a**, Fluorescence response (z-scored Δ*F/F*_0_) along time and trials during reception of a *CS^−^* odor cue (1*s*, top panels) and during learning of the association between a different odor, *CS*^+^ (1*s*), and a fixed size reward delivery presented 3*s* after the odor initiation (bottom panels) across NAc and DMS regions. **b**, Quantification of *CS*^+^ response across trials (top), of outcome response across trials (*US*, middle), and of temporal position of the peak activity during the delay period (i.e. from cue initiation to reward delivery) across trials. Lines represent a linear fit of each one of these variables to trial number. Slopes (*β*) and corresponding significance are shown for each striatal region of this example mouse. **c**, same as **a** for DLS and TS regions. **d**, same as **b** for DLS and TS regions.

**Extended Data Figure 6.**
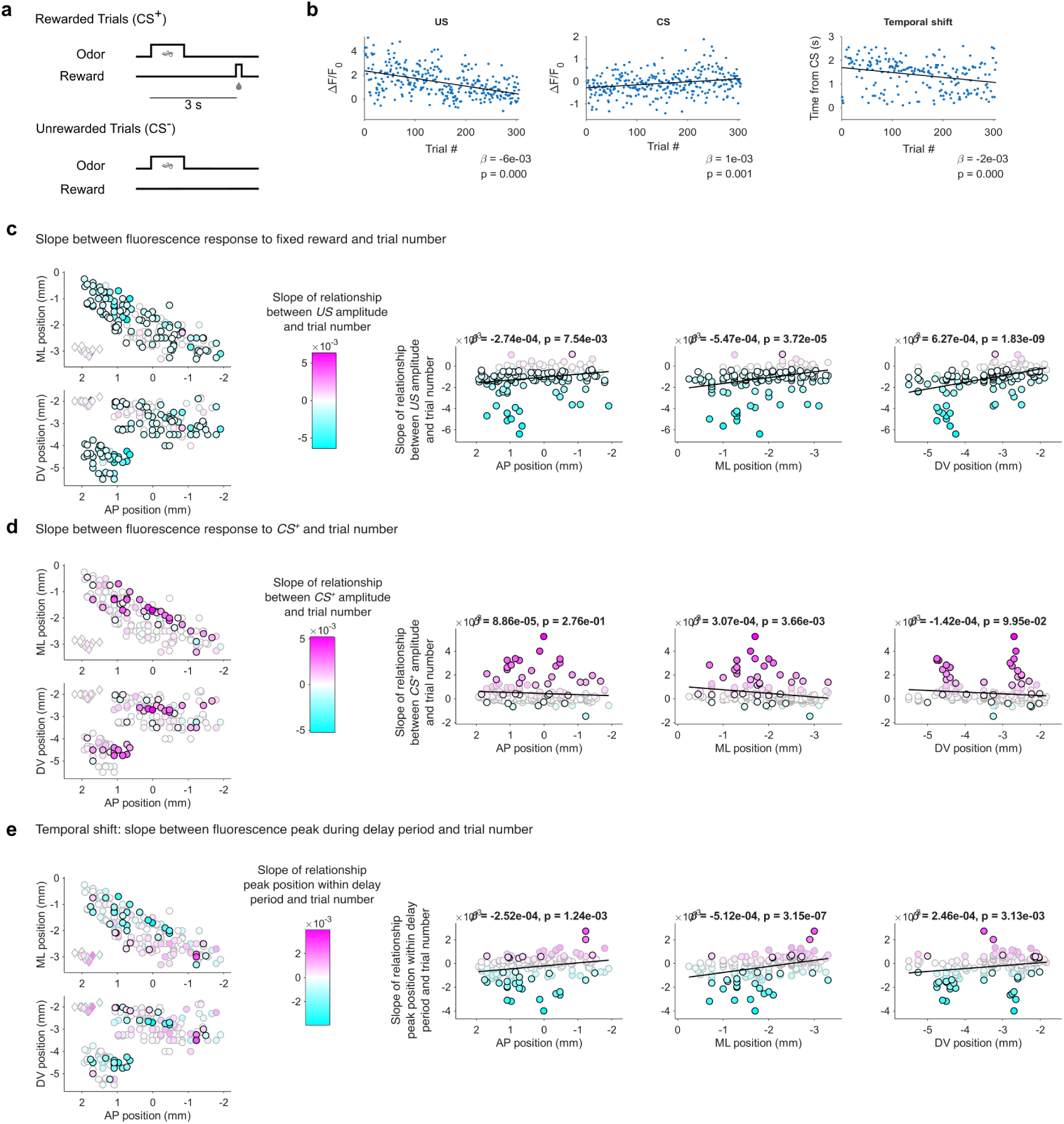
Modulation of cue and outcome responses during learning across striatal regions. **a**, Classical conditioning task used to quantify modulation of cue, outcome and temporal shift activity during the learning process. **b**, Example modulation of outcome (*US*, left), cue (*CS*^+^, middle), and temporal position of peak activity during the delay period (from cue initiation to reward delivery) across trials from the lNAc of example mouse shown in Extended Data Fig. 5. Slopes (*β*) and corresponding significance of linear fit with trial number were obtained for all fibers. **c**, Slope between fluorescence response (z-scored Δ*F/F*_0_) to the fixed reward delivery and trial number (14 fibers for each mouse, *n* = 11 mice). Left: data shown in the AP-ML projection (top, horizonal view) and AP-DV projection (bottom, sagittal view). Diamonds plotted around the (2*AP, −*3*ML, −*2*DV*) coordinates correspond the control ROIs in the cameras’ fields of view, not to real fibers in the brain. Fibers with black edge show significant modulation with trial number (*P <* 0.05 for slope *β*), fibers with grey edge signal non-significant modulation with trial number. Right: Same data shown in each anatomical axis with slope (*β*) and significance value of a linear fit of the data to each anatomical axis. **d**, Same as **c** for the modulation of *CS*^+^ with learning. **e**, Same as **c** for the temporal position of peak activity during the delay period over trials.

**Extended Data Figure 7.**
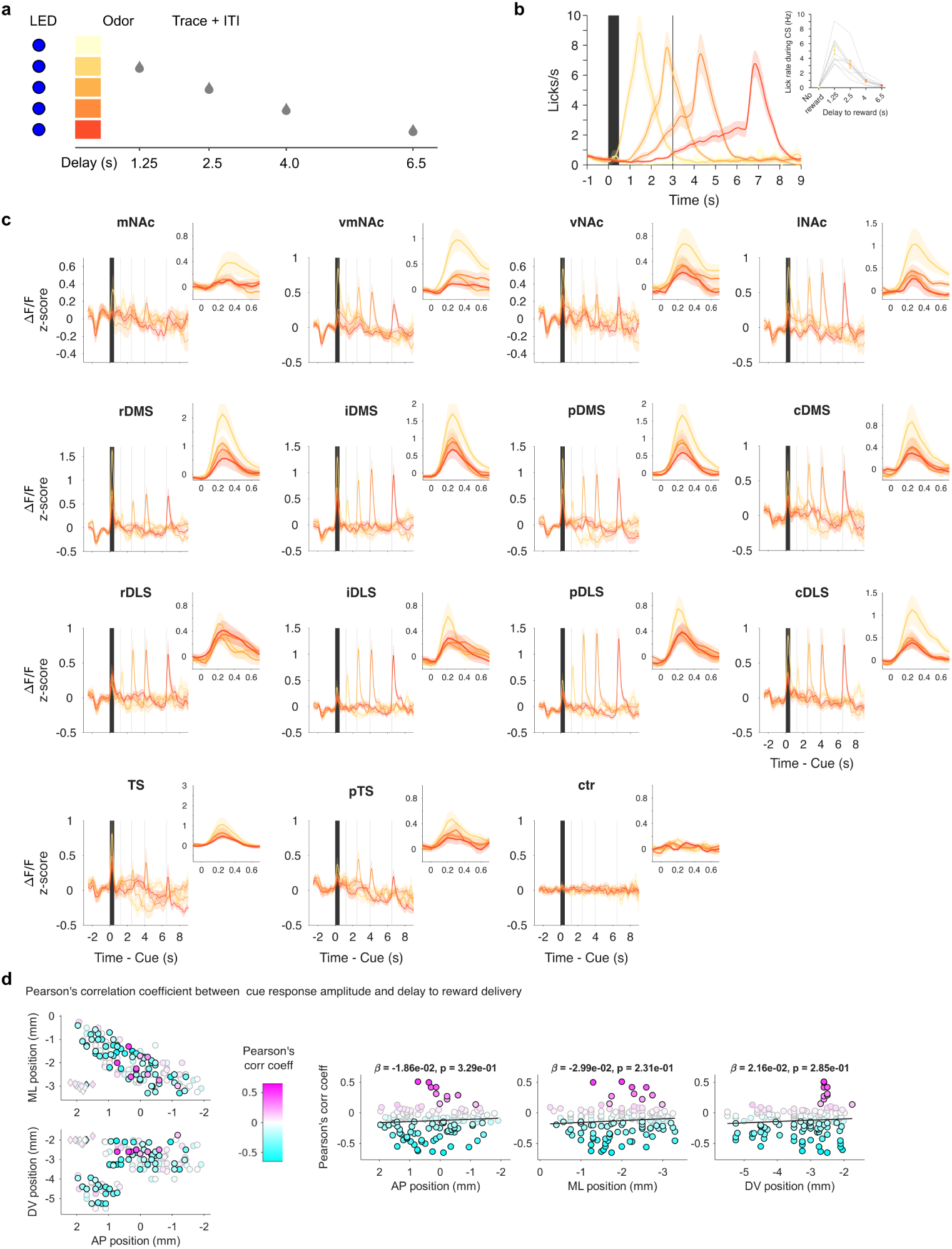
Mapping of temporal discounting during cue activity across the striatum. **a**, Classical conditioning task to evaluate temporal discounting of cue activity across striatal regions: 4 different odors predict reward delivery following delays of 1.25, 2.5, 4 and 6.5*s*. **b**, Average licking pattern for each one of the rewarded odor cues. Inset: mean lick rate during the *CS*^+^ (lines represent individual mice, *n* = 10). **c**, Trial-averaged fluorescence (z-scored Δ*F/F*_0_) along trial time for the 4 rewarded trial types across striatal targets. Insets: Zoom-in view of activity during the cue presentation (0 *−* 0.5*s*). **d**, Pearson’s correlation coefficient between fluorescence response (z-scored Δ*F/F*_0_) to the odors cues and corresponding delay to reward delivery (14 fibers for each mouse *n* = 10 mice). Left: data shown in the AP-ML projection (top, horizonal view) and AP-DV projection (bottom, sagittal view). Diamonds plotted around the (2*AP, −*3*ML, −*2*DV*) coordinates correspond the control ROIs in the cameras’ fields of view, not to real fibers in the brain. Fibers with black edge show significant correlation (*P <* 0.05), fibers with grey edge show non-significant correlations. Right: Same data shown in each anatomical axis with slope (*β*) and significance value of a linear fit of the data to each anatomical axis.

**Extended Data Figure 8.**
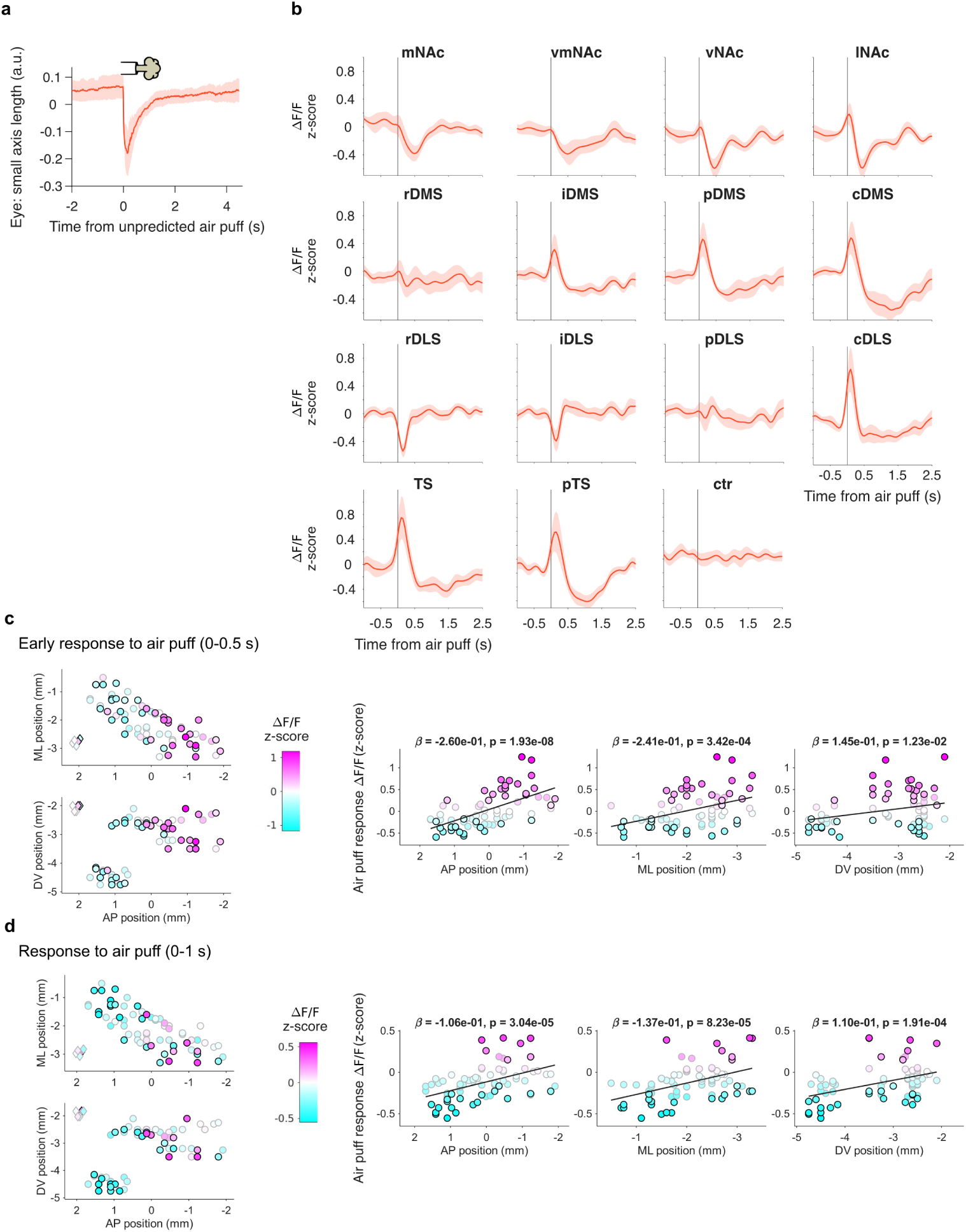
Mapping responses to air puff across the striatum. **a**, Quantification of the length of the smaller axis of the eye during air puff delivery. **b**, Fluorescence response (z-scored Δ*F/F*_0_) to the air puff delivery across striatal targets and averaged across mice (*n* = 5). **c**, Average fluorescence response (z-scored Δ*F/F*_0_, 0 *−* 0.5*s*) to air puff delivery measured in each fiber (14 fibers in each of *n* = 5 mice). Left: data shown in the AP-ML projection (top, horizonal view) and in the AP-DV projection (bottom, sagittal view). Diamonds plotted around the (2*AP, −*3*ML, −*2*DV*) coordinates correspond the control ROIs in the cameras’ fields of view, not to real fibers in the brain. Fibers with black edge show significant modulation from baseline (*P <* 0.05, Wilcoxon sign rank test), fibers with grey edge signal non-significant modulation from baseline. Right: Same data shown in each anatomical axis with slope and significance value of a linear fit of the data to each anatomical axis. **d**, same as **c** for the 0 *−* 1*s* period.

**Extended Data Figure 9.**
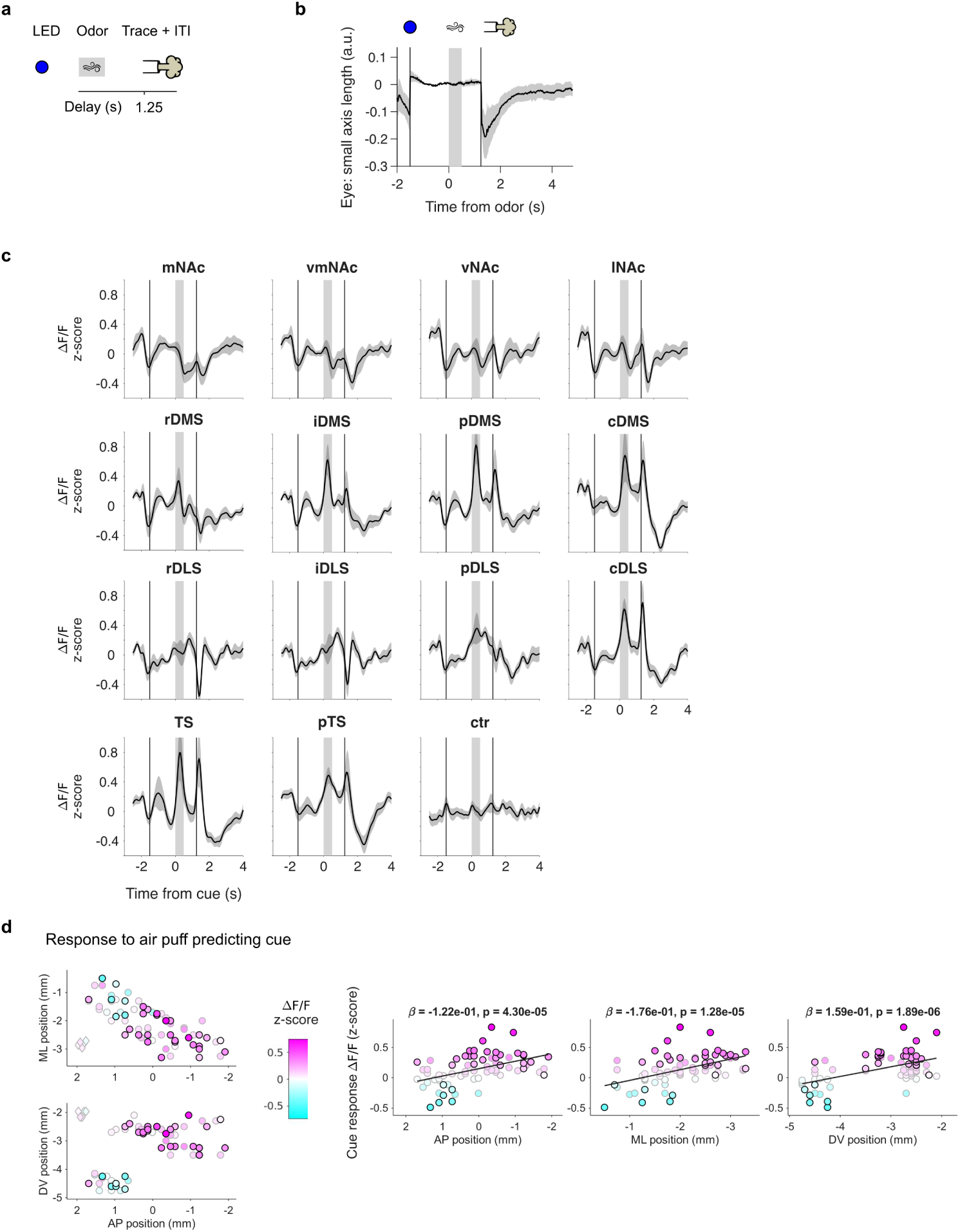
Mapping responses to air puff predicting cue across the striatum. **a**, Temporal sequence of events leading to air puff delivery. **b**, Average length of the smaller axis of the eye during the trials. **c**, Trial-averaged fluorescence (z-scored Δ*F/F*_0_) along trial time across striatal targets. **d**, Average fluorescence response (z-scored Δ*F/F*_0_) to air puff predicting cue measured in each fiber (14 fibers in each of *n* = 5 mice). Left: data shown in the AP-ML projection (top, horizonal view) and in the AP-DV projection (bottom, sagittal view). Diamonds plotted around the (2*AP, −*3*ML, −*2*DV*) coordinates correspond the control ROIs in the cameras’ fields of view, not to real fibers in the brain. Fibers with black edge show significant modulation from baseline (*P <* 0.05 Wilcoxon sign rank test), fibers with grey edge signal non-significant modulation from baseline. Right: Same data shown in each anatomical axis with slope and significance value of a linear fit of the data to each anatomical axis.

**Extended Data Figure 10.**
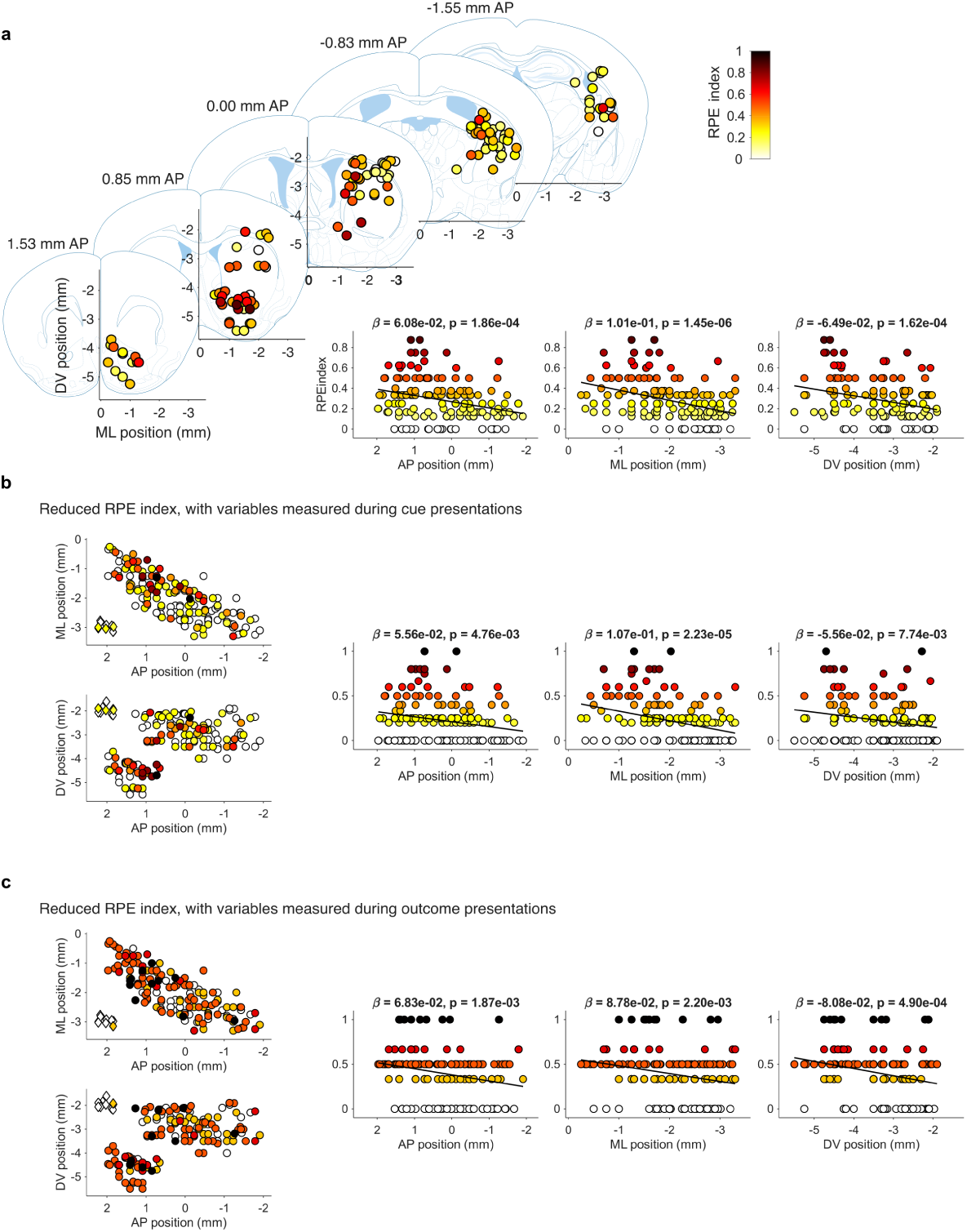
Anatomical gradients for canonical RPE index across the striatum. **a**, RPE index colapsed to 5 coronal brain slices (top left, same slices used in Fig. 2) and as a function of fiber location along the anterior-posterior, medio-lateral, and dorso-ventral axis of the stiatum (bottom right), with corresponding slope (*β*) of linear fit to axis coordinate and significance of the slope. **b** Same as **a** considering only the criteria that rely on RPE signal during cue presentation. **c** Same as **a** considering only the criteria that rely on RPE signal during outcome presentation.

**Extended Data Figure 11.**
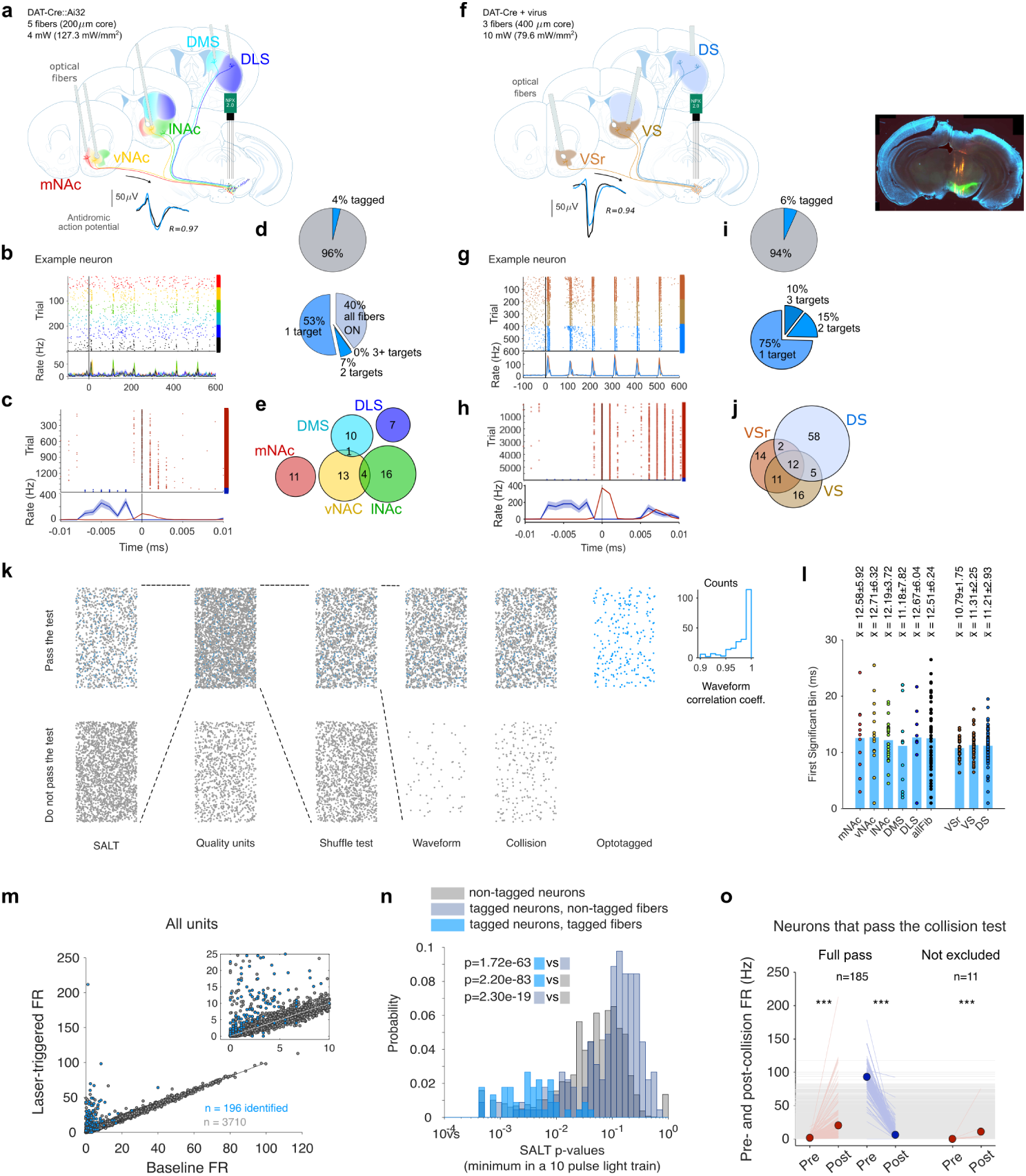
Experimental strategy for acute recording and identification of projection-defined dopamine neurons. **a**, Experimental strategy for antidromic identification of midbrain dopamine neurons using a custom five-fiber implant for photostimulation through each fiber. Example spontaneous (black) and antidromically (blue) evoked spike waveforms are shown below the schematic together with the correlation coefficient between them. **b**, Example raster plot and peri-stimulus time histogram (PSTH) showing laser-evoked antidromic responses for a representative neuron. Colored rasters denote different stimulated projection targets as in the schematic in **a** (in black trials all fibers where illuminated simultaneously, allowing the identification of a larger number of dopamine neurons, but for which we do not know the projection target). **c**, Collision test for the example neuron shown in **a,b**. Spike trains were aligned to the first significant antidromic response bin following laser stimulation. Spontaneous spikes occurring within the expected antidromic latency (*−*7 to *−*2*ms* relative to antidromic spike onset) abolished the laser-evoked spike, confirming antidromic activation. **d**, Summary of optotagging yield for the five-fiber experimental configuration. Top: proportion of recorded neurons classified as optotagged. Bottom: fraction of neurons projecting to one, two, three, or more stimulation targets. **e**, Overlap between projection-defined populations identified in the five-fiber experiment. Numbers indicate neurons antidromically identified from each projection target combination. **f**, **g**, **h**,**i**, and**j**, Same as **a**, **b**, **c**, **d**, and **e**, respectively, for the three-fiber configuration. **k**, Optotagging pipeline used to identify projection-defined dopamine neurons. Neurons were required to pass unit-quality metrics, a shuffle test for laser responsiveness, waveform similarity between spontaneous and laser-evoked spikes, a collision test, and the Stimulus-Associated spike Latency Test (SALT). Blue dots in the top tow indicate neurons passing all criteria, grey dots indicate all the other units. **l**, Latency of the first significant laser-evoked response (in relation to a shuffled control) across projection targets. Points represent individual projection-specific neuron–target pairs; bars indicate the mean. **m**, Mean firing rate during the laser response as a function of baseline firing rate for all recorded units (gray) and optotagged neurons (blue). Inset shows the low firing-rate range. **n**, Distribution of SALT p-values for non-optotagged neurons (gray), for optotagged neurons given stimulation sites that did not antidromically identify the neuron (dark blue), and for optotagged neurons given projection targets satisfying all optotagging criteria (light blue). P-values shown with shaded squares correspond to the comparison of the different distributions using two-sample Kolmogorov-Smirnov test. **o**, Validation of the collision test: mean firing rates immediately before and after the expected antidromic spike are shown separately for neurons fully passing the collision test and for neurons not excluded because of no spontaneous spikes within the collision window. Thin lines represent individual neurons; circles denote population me_5_an_2_s. *^∗∗∗^P <* 0.001 (Wilcoxon signed-rank test).

**Extended Data Figure 12.**
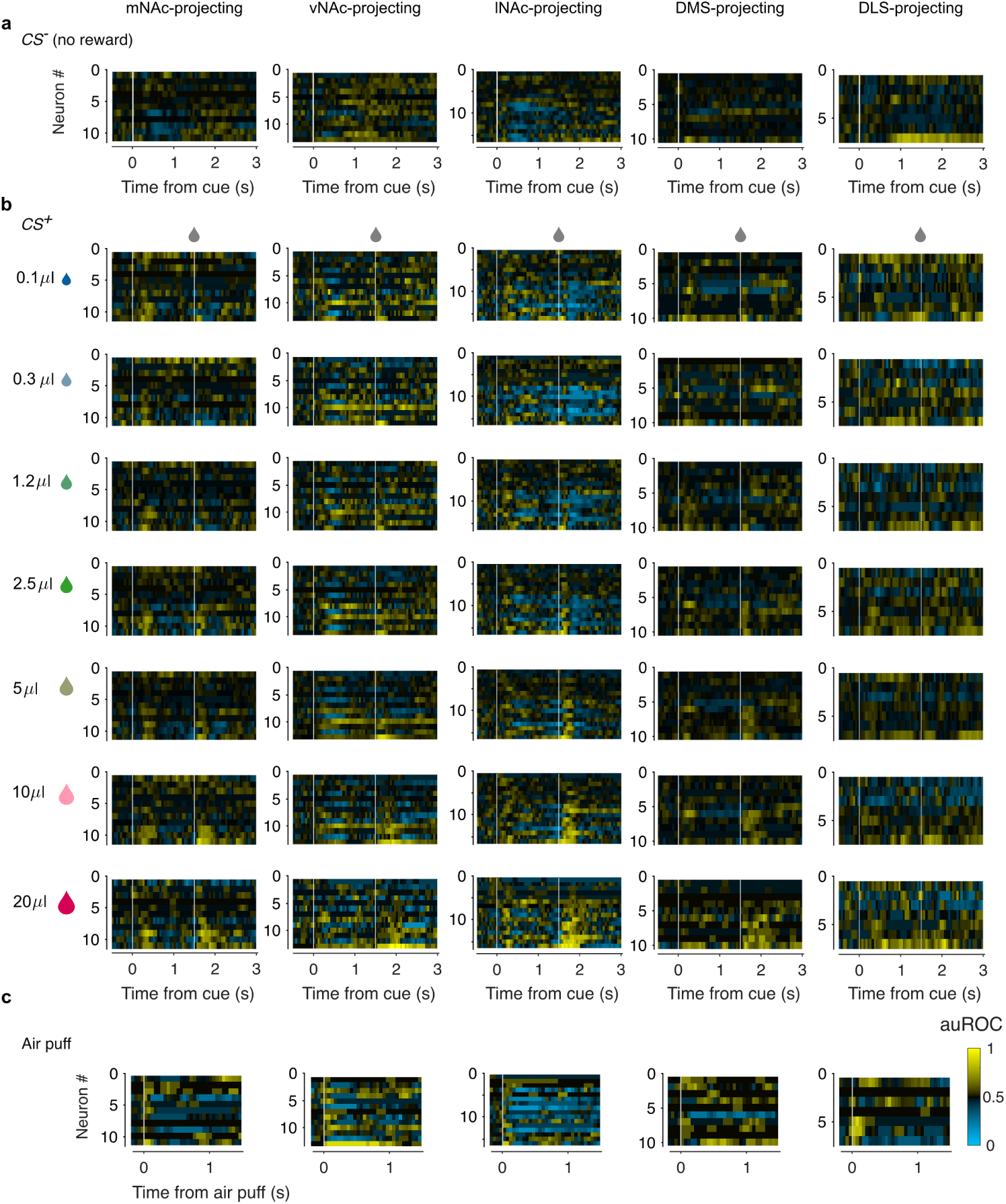
Projection-defined dopamine neuron activity. **a**, Mean activity of individual dopamine neurons during *CS^−^* trials, separated by projection target (i.e. optotagging label). Activity is represented as area under the receiver-operator curve in 50*ms* bins and considering baseline the last second before cue presentation. **b**, Same as **a** for *CS*^+^ trials: each row shows the mean activity during trials in which a particular reward size was delivered. **c**, Same as **a** for air puff trials.

**Extended Data Figure 13.**
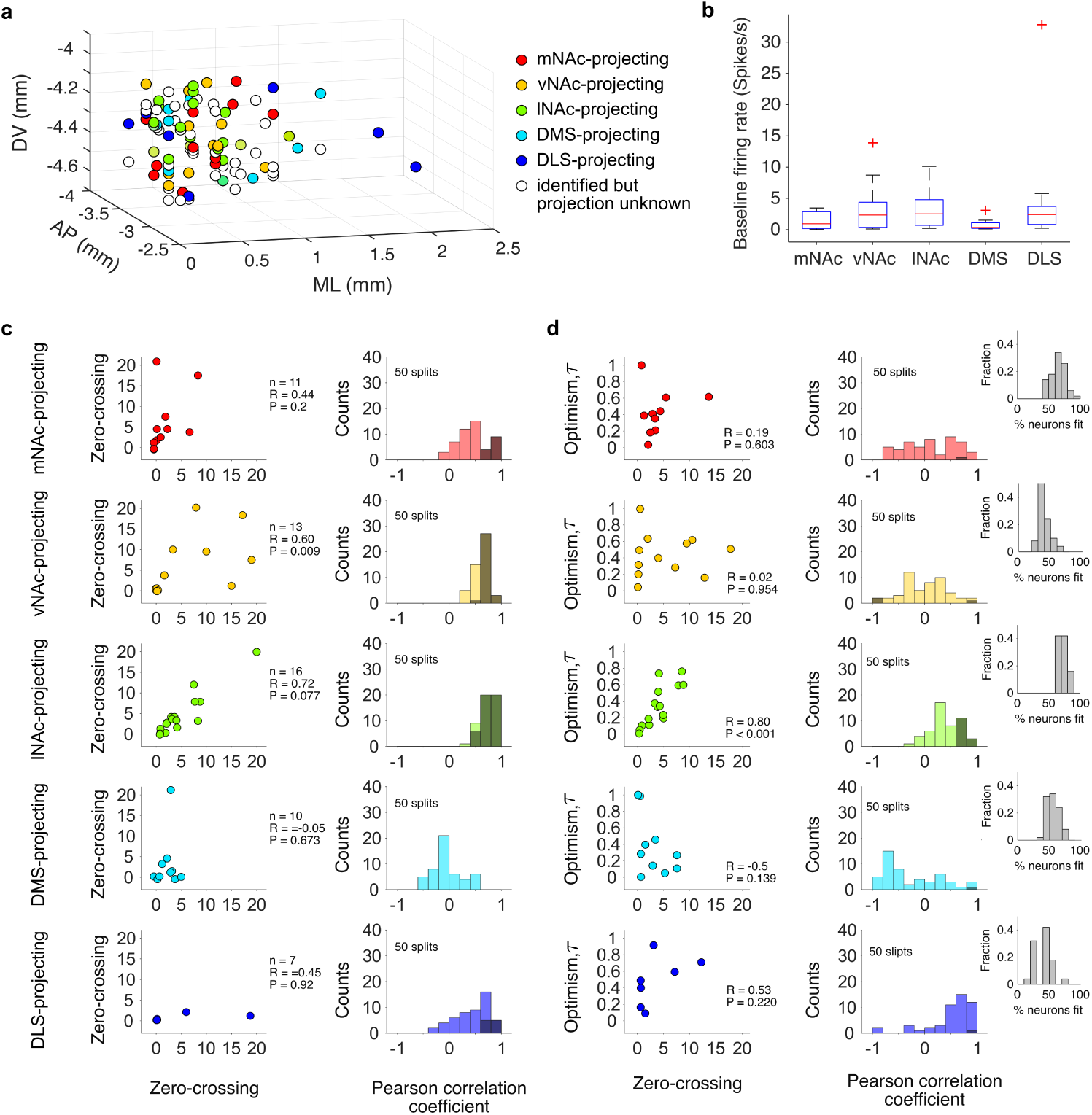
Empirical calculation of zero-crossing point and optimism level across projectiondefined dopamine populations. **a**, Anatomical location of dopamine neurons recorded in mice implanted with the custom 5-fiber implant. Neuros are color-coded based on their projection target. **b**, Baseline firing rate of recorded dopamine neurons per projection target. **c**, Left: Relationship between zero-crossing points calculated in two separate halves of the data (1 split) for dopamine neurons projecting to different striatal targets (rows). Right: histogram of Pearson’s correlation coefficients obtained across 50 splits of the data, significant splits are shown in darker shade. **d**, Left: Average optimism level as a function of average zero-crossing point for each projection-defined population across 50 splits of the data in two halves. Right: histogram of Pearson’s correlation coefficients obtained for the 50 splits of the data, significant splits are shown in darker shade. Insets show the histogram of the percentage of neurons successfully fit across the 50 splits.

**Extended Data Figure 14.**
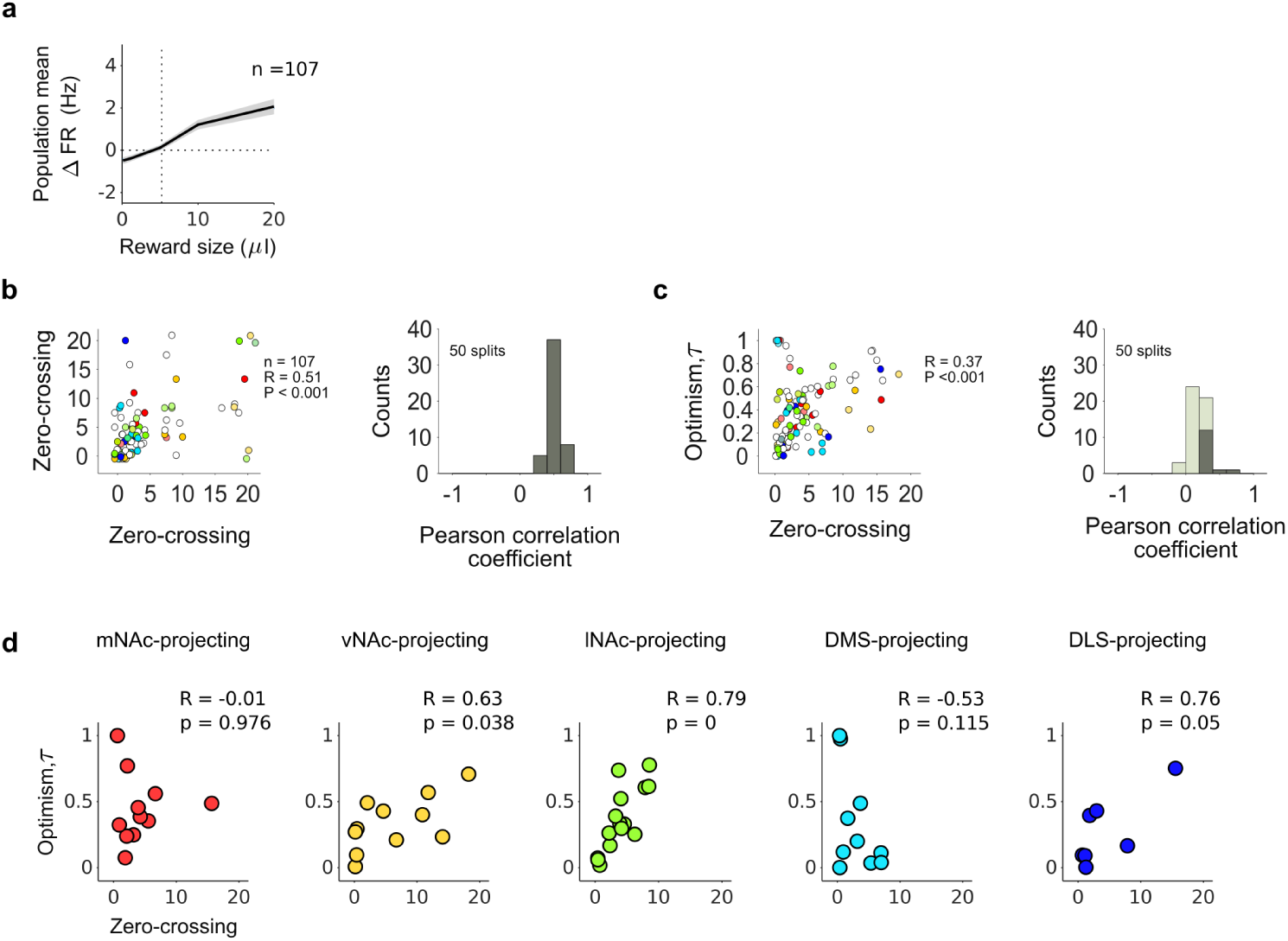
Empirical calculation of zero-crossing point and optimism after pooling all dopamine neurons (instead of separating them by projection target). **a**, Empirical calculation of zero-crossing point and optimism without splitting the data into different projection-defined populations, i.e., pooling all identified dopamine neurons together to obtain their global response function to reward size (instead of a response function per projection target, as in Fig. 3 and Extended Data Fig. 13). **b**, Relationship between zero-crossing points calculated in two separate halves of the data (1 split) for all dopamine neurons pooled together. Right: histogram of Pearson’s correlation coefficients obtained across 50 splits of the data, all splits were significant and are shown in darker shade. **c**, Left: Average optimism level as a function of average zero-crossing point for all recorded dopamine neurons across 50 splits of the data in two halves. Right: histogram of Pearson’s correlation coefficients obtained for the 50 splits of the data, significant splits are shown in darker shade. **d**, Post-analysis separation of the data shown in **c** into the different striatal projection targets: when all neurons are pooled together to obtain a global response function for converting zero-crossing point from neural activity to reward size (as in **a**) variability across populations is reduced, revealing more neurons that could be contributing to the outcome representation in all projection targets. Still, lNAc neurons show the strongest and most significant correlation between these two variables.

**Extended Data Figure 15.**
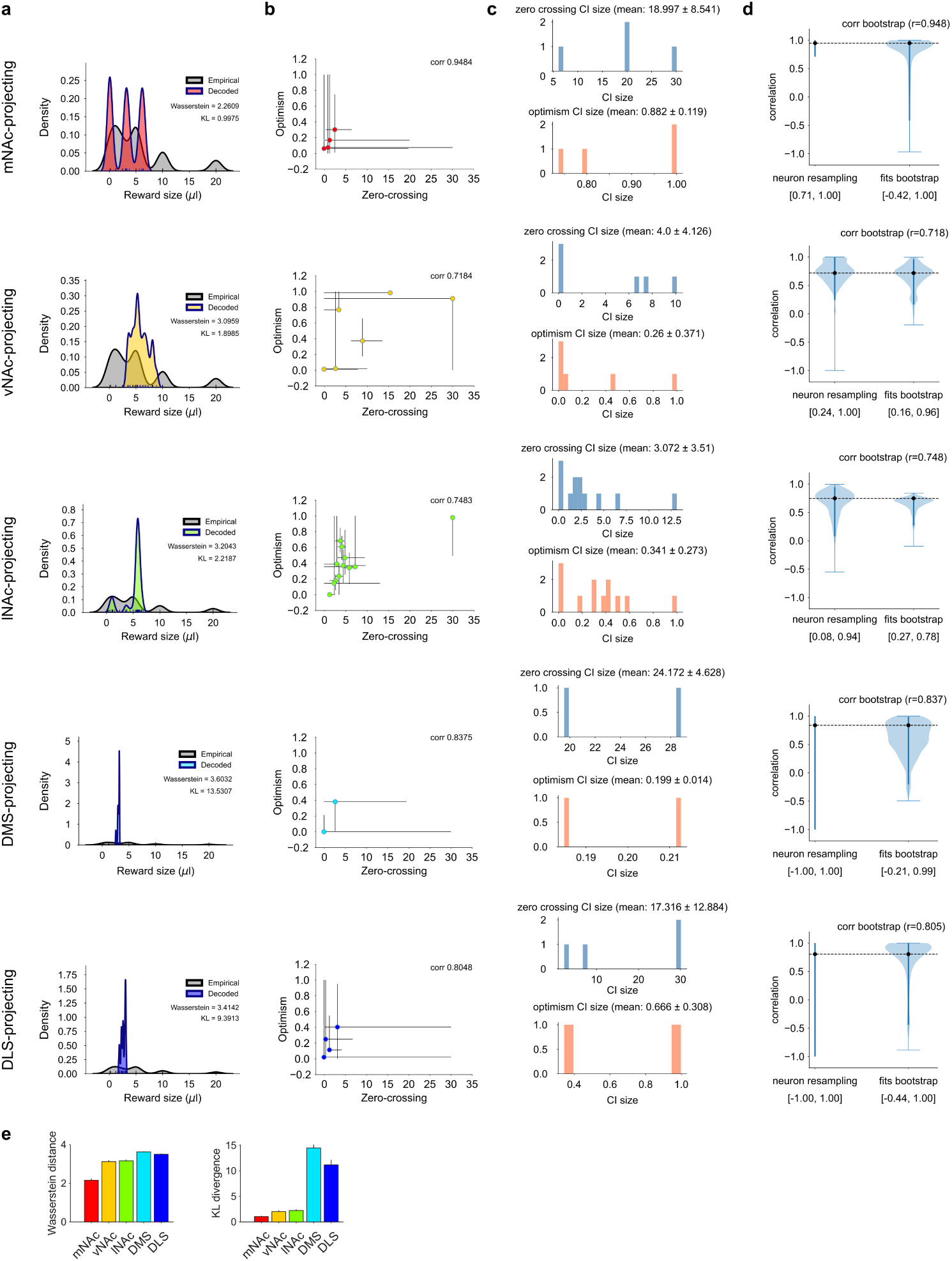
Model-based estimation of zero-crossing point and optimism level of projection-defined dopamine neurons. For each projection-target (rows): **a**, Reward distribution decoded with the expectile decoder from the MLE estimates of zero-crossing and optimism of a projection-defined population (colored) overlaid on the true reward distribution (grey), with Wasserstein distance and KL divergence shown in the inset text. **b**, Optimism versus zero-crossing point for the fitted neurons of each projection-defined population (as in Fig. 3k), with 95% confidence intervals from a non-parametric bootstrap over trials. **c**, Distribution of confidence-interval widths for zero-crossing (top) and optimism (bottom). **d**, Bootstrap distributions of the Pearson correlation between zero-crossing and optimism, obtained by resampling neurons (left violin) or by recomputing the correlation across neurons on each bootstrap refit (right violin); dashed line, point estimate; brackets, 95% CI. **e**, Wasserstein distance (left) and KL divergence (right) between decoded and true distributions, mean *±* SD over 10 decoding runs.

**Extended Data Figure 16.**
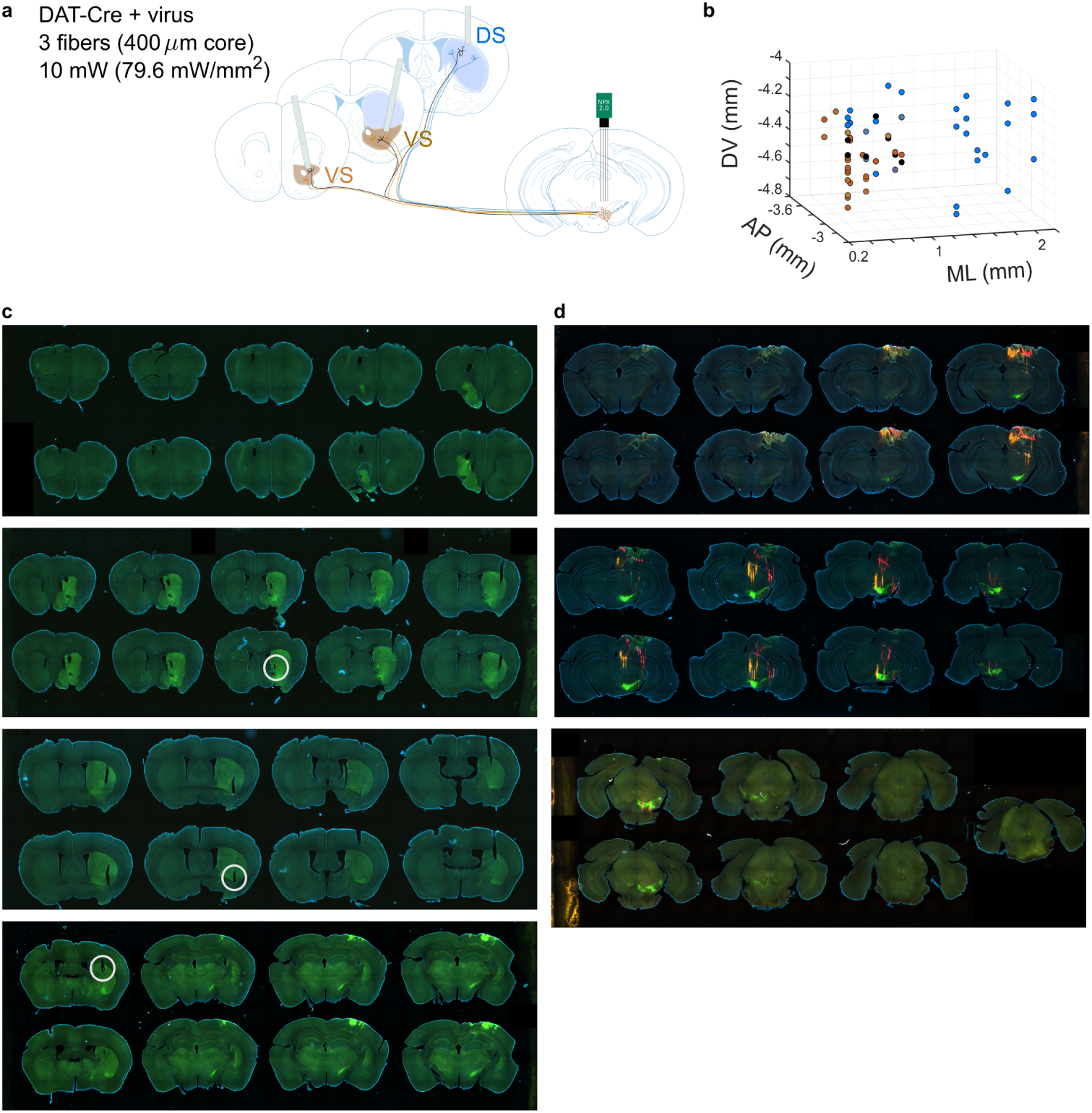
Projection-defined recording of dopamine neuron activity in a single mouse. **a**, Schematic of planned experiment to uncover distribution information in a single mouse. **b**, Anatomical location of the recorded neurons in the VTA and SNc: AP: anterior-posterior position, ML: medial-lateral position, DV: dorso-ventral position. Colors correspond to projection targets in the striatum as in **a**: ventral striatum in brown and dorsal striatum in blue. **c**, Histological sections of the striatum of this mouse, the location of the tips of the optical fibers are circled in white. **d**, Histological sections showing transfected dopamine neurons in green and shanks of the Neuropixels 2.0 probes in red (DiD) and orange (DiI).

**Extended Data Figure 17.**
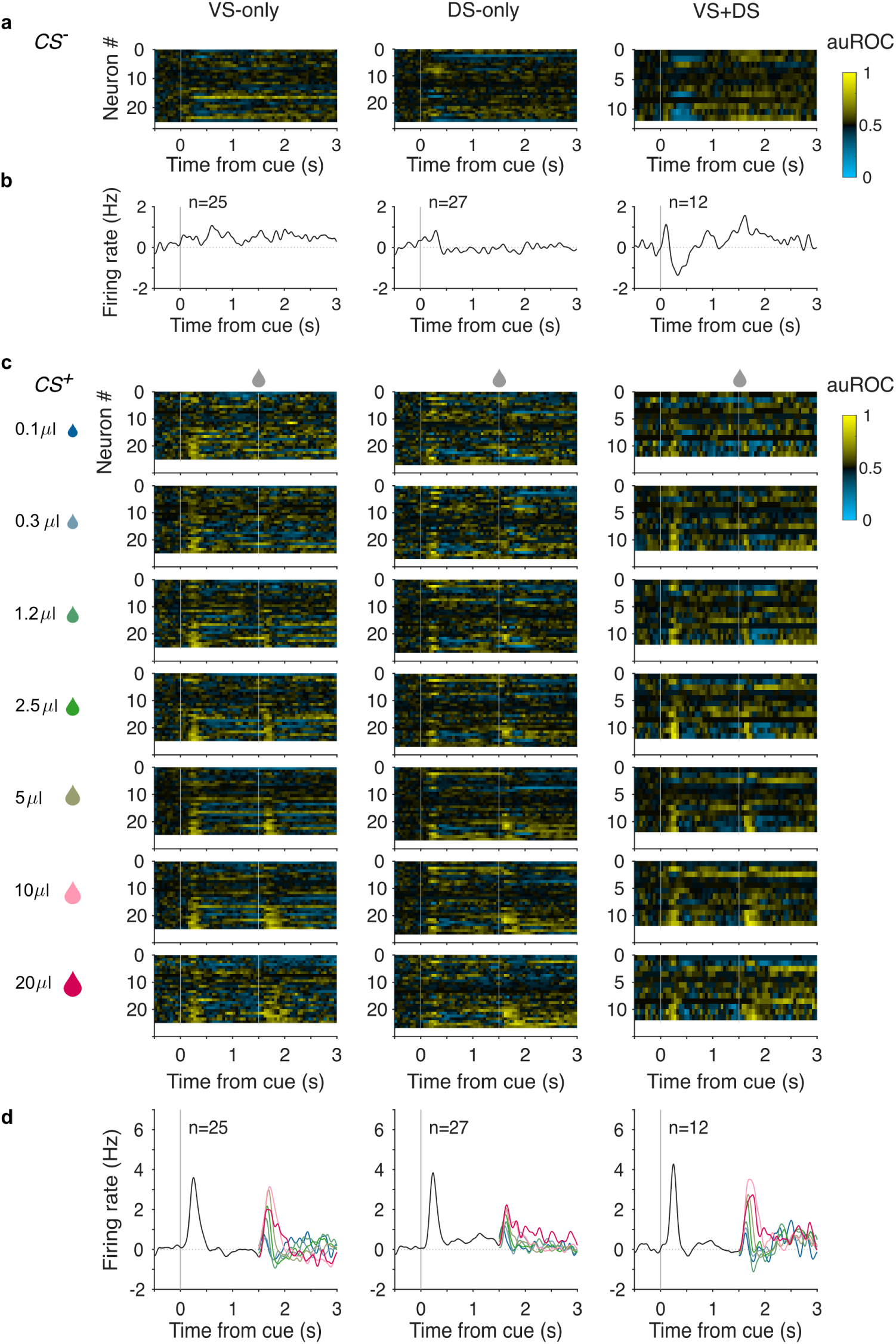
Projection-defined dopamine neuron activity. **a**, Mean activity of individual dopamine neurons during *CS^−^* trials, separated by projection target (i.e. optotagging label). Activity is represented as area under the receiver-operator curve in 50*ms* bins and considering baseline the last second before cue presentation. **b**, Population mean firing rate to *CS^−^* trials. **c**, Same as **a** for *CS*^+^ trials: each row shows the mean activity during trials in which a particular reward size was delivered. **d**, Population average firing rate during *CS*^+^ trials: colors following the reward delivery represent the different reward sizes.

**Extended Data Figure 18.**
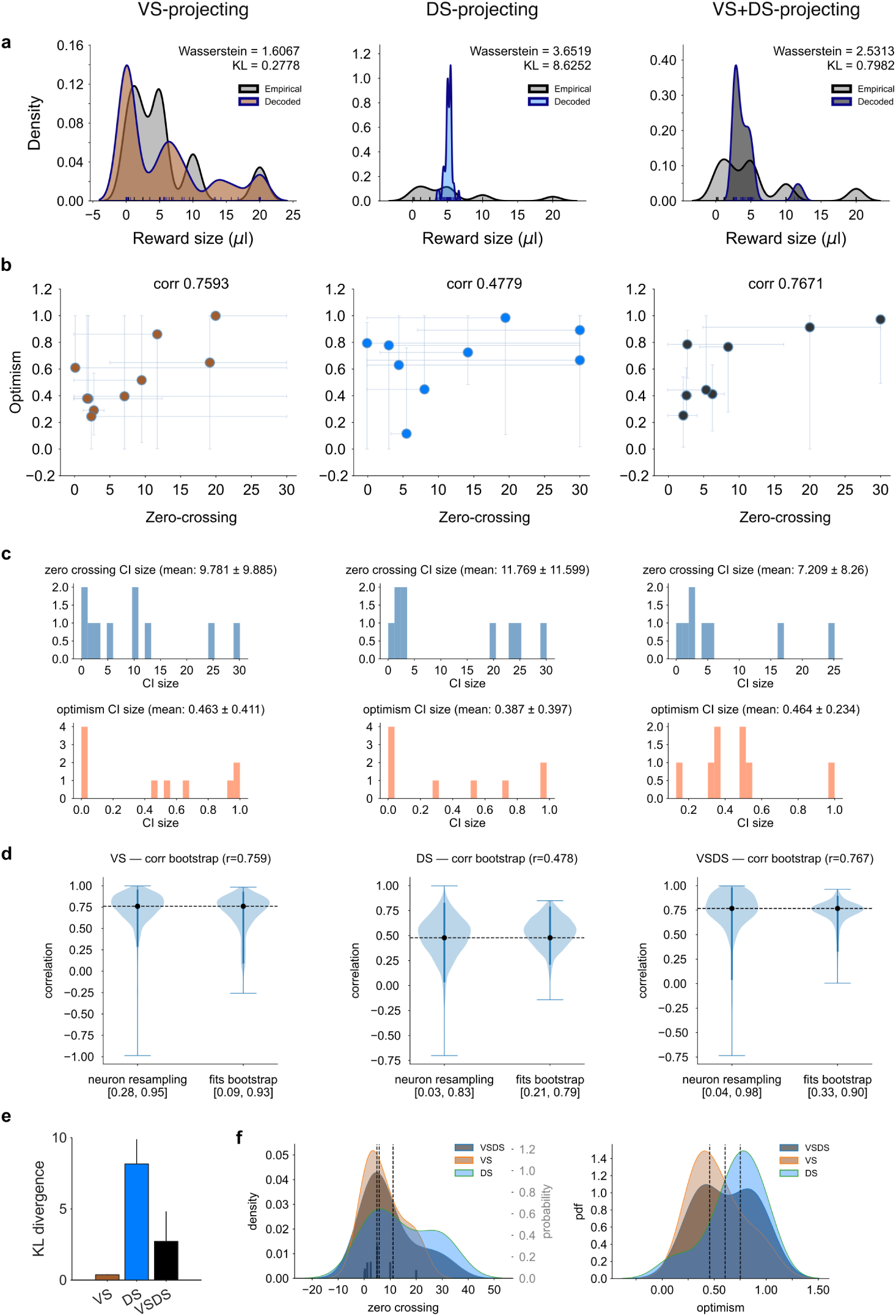
Model-based estimation of zero-crossing point and optimism level of dopamine neurons recorded in a single mouse. **a**, Reward distribution decoded with the expectile decoder from the MLE estimates of zero-crossing and optimism of each population (blue) overlaid on the true reward distribution (grey), with Wasserstein distance and KL divergence shown in the inset text. **b**, Optimism versus zero-crossing point for each fitted neuron (as in Fig. 4B), with 95% confidence intervals from a non-parametric bootstrap over trials. **c**, Distribution of confidence-interval widths for zero-crossing (top) and optimism (bottom). **d**, Bootstrap distributions of the Pearson’s correlation between zero-crossing and optimism, obtained by resampling neurons (left violin) or by recomputing the correlation across neurons on each bootstrap refit (right violin); dashed line, point estimate; brackets, 95% CI. **e**, KL divergence between decoded and true distributions (mean *±* SD over 10 decoding runs). **f**, Kernel density estimates of zero-crossing (left) and optimism (right) for each population; dashed lines, medians; true reward distribution shown as grey bars.

**Extended Data Figure 19.**
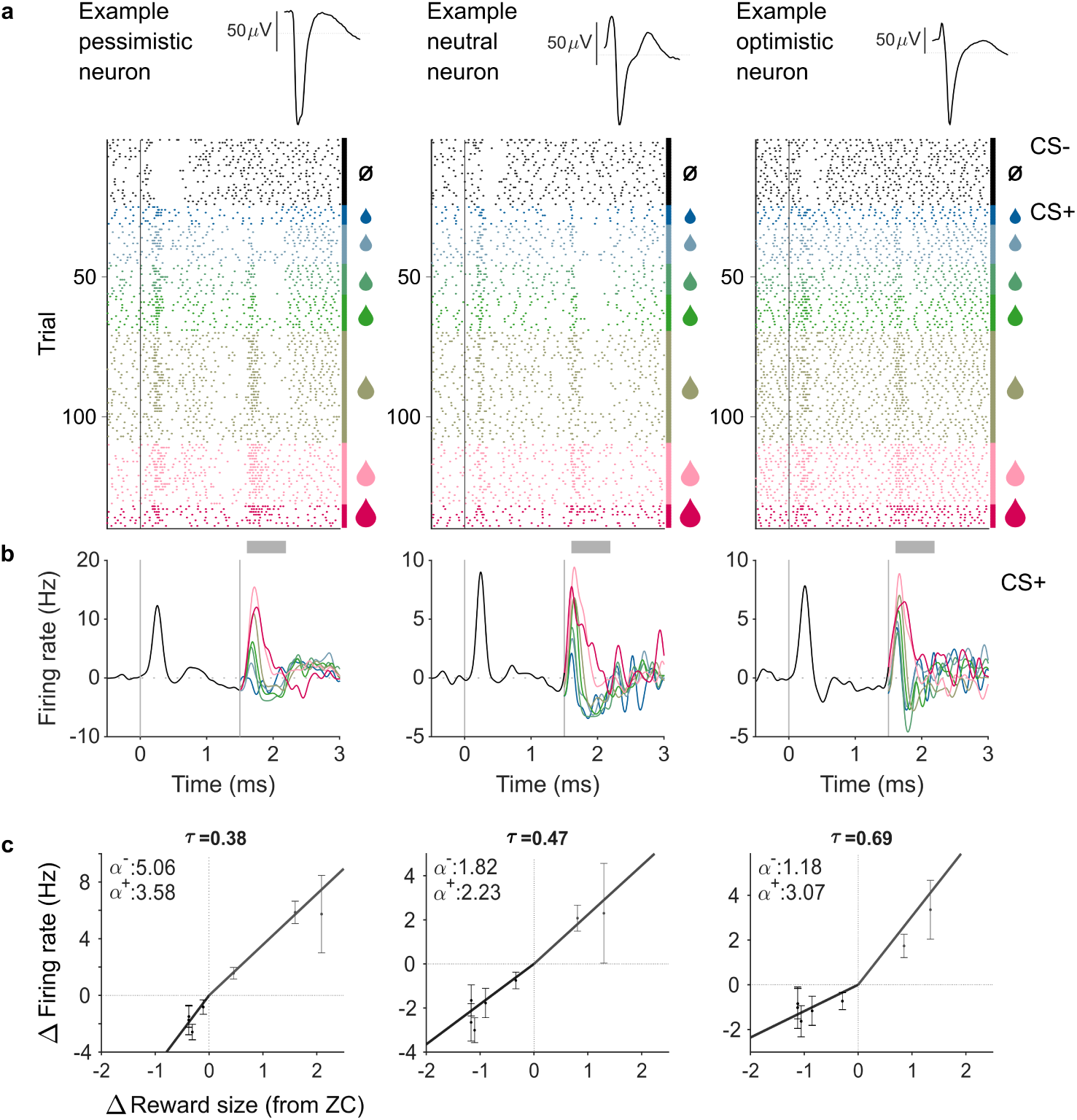
Example pessimistic, neutral, and optimistic neurons. **a**, Spike waveforms and raster plots showing the action potential timings of three example neurons during the task. For display purposes, trials are grouped into *CS^−^* (black) and *CS*^+^ and in ascending order for reward size (but are interleaved during task performance). **b**, Mean single neuron activity for *CS*^+^ trials (color after reward delivery time corresponds to reward size; gray bar represents the window of activity used for further analysis, such as zero-crossing estimation). **c**, Change in firing rate, quantified in the period 200 *−* 600*ms* after reward delivery (gray bar in **b**) as a function of the difference between delivered reward size and zero-crossing point. Zero-crossing point for each cell is defined as the origin of the plots, corresponding to the reward size that elicits no prediction error.

**Extended Data Figure 20.**
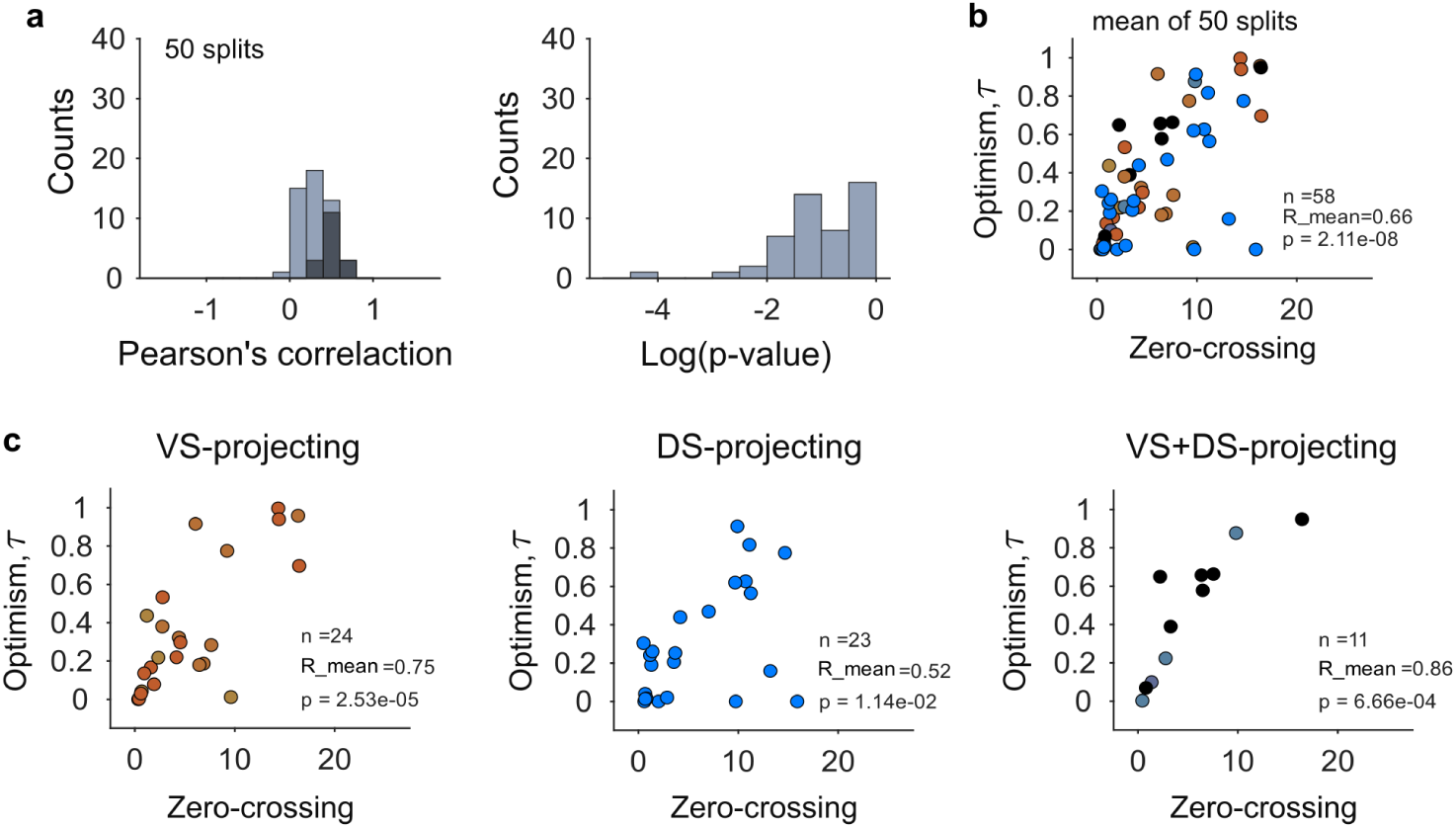
Empirical calculation of zero-crossing point and optimism level of dopamine neurons recorded in a single mouse. **a**, Trial data was split into two halves for empirical calculation of zero-crossing point and optimism, a procedure that was repeated 50 times for evaluating the relationship between zero-crossing and optimism level (*τ*) (*n* = 58 neurons). Neurons for which this procedure fails (*n* = 6, 1 from VS, 4 from DS, 1 from VS+DS) are not used for further analysis. Left: histogram of correlation coefficients obtained across 50 splits (significant correlation coefficients are shown in dark gray). Right: histogram of p-values for the correlation coefficients obtained across those 50 splits. **b**, Average optimism level as a function of average zero-crossing point for these 58 cells. **c**, Same data as in **b**, with cells separated based on their projection target (in the VS+DS plot black dots show neurons tagged by all three fibers and grey dots represent neurons tagged by one VS fiber and the DS fiber). A significant positive correlation is observed across the three populations, the weakest of which is seen for DS-projecting dopamine neurons, which deviate considerably from the expectile prediction (see main figure 4). The population that best follows the expectile prediction is the VS+DS population.

**Extended Data Figure 21.**
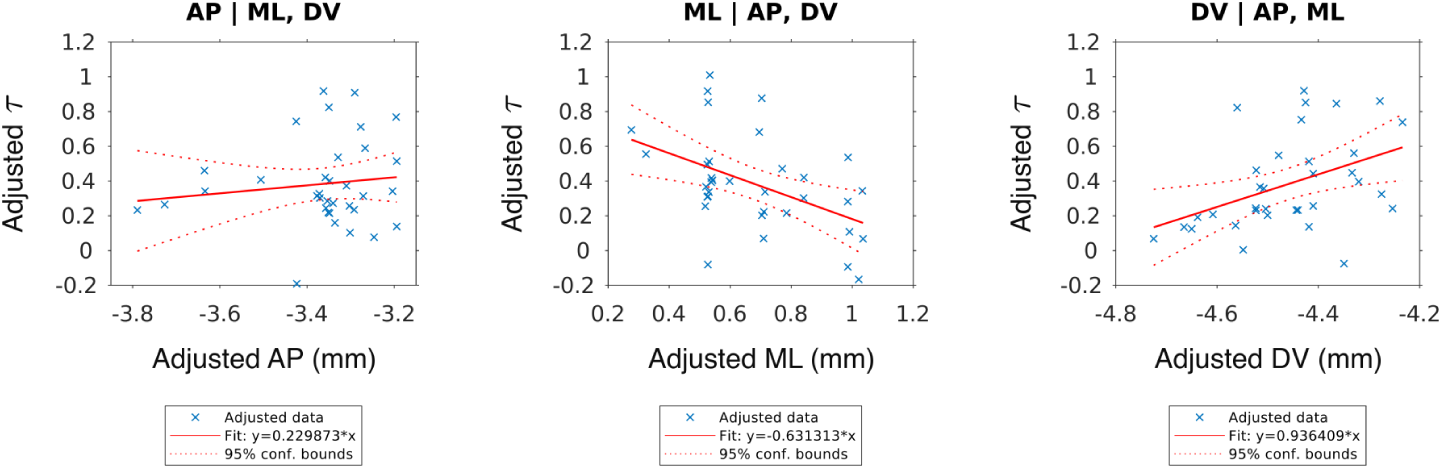
Anatomical organization of dopamine neuron optimism after accounting for three-dimensional location. Partial regression plots showing the relationship between optimism (*τ*) and each anatomical coordinate (AP, ML, DV), controlling for the other two coordinates. Multiple linear regression was performed on 34 VS- and VS+DS-projecting dopamine neurons recorded from one mouse. The overall model was significant (*R*^2^ = 0.396, *F*_3,30_ = 6.56, *P* = 0.0015). Optimism was negatively associated with ML position (*β* = *−*0.631, *P* = 0.0065) and positively associated with DV position (*β* = 0.936, *P* = 0.0158), but not significantly associated with AP position (*β* = 0.230, *P* = 0.475). Coefficients and P values correspond to the multiple regression model; each cross represents one neuron. Correlation analysis on the neurons’ coordinates reveals that correlations are weak (AP-ML: −0.498, AP-DV: −0.109, ML-DV: 0.081), excluding confounds due to collinearity among the predictors.

**Extended Data Figure 22.**
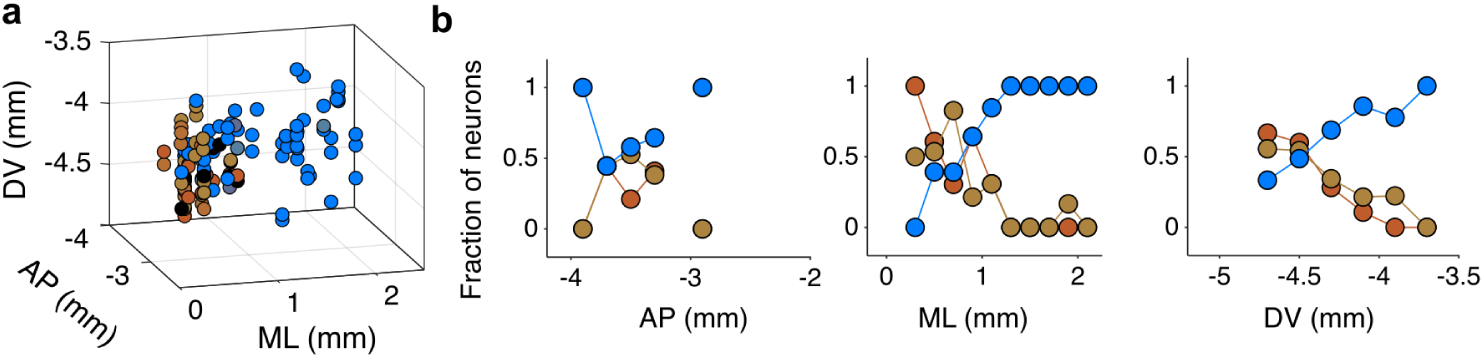
Projection targets of the recorded midbrain dopamine neurons. **a**, Anatomical location of all dopamine neurons recorded in a preparation of two fibers targeting lateral VS and one targeting DS, pooled across 5 mice, color coded according to projection target (brown: VS fibers, blue: DS fiber). **d**, For each anatomical bin of 200*µm* in the midbrain. we plot the fraction of neurons projecting to each fiber location (color) in the AP (left), ML (center), and DV (right) axis. Logistic regression applied on the three axis results in a significant anatomical organization in the ML and DV axis for the most rostral lateral VS fiber (*P <* 0.001 ML, DV, n.s. AP), a significant ML organization for the posterior lateral VS fiber (*P <* 0.001 ML, n.s. AP and DV), and a significant organization in all three anatomical axis for the DS fiber (*P* = 0.011 AP, *P <* 0.001 ML, *P* = 0.008 DV).

